# Engineering a Novel Bacterial Encapsulin for Programmable Surface Functionalization: From Single-Target to Mosaic Nanovaccines

**DOI:** 10.64898/2026.06.01.729406

**Authors:** India Boyton, Claire Rennie, Julia van der Hoven, Manuel Del Valle, Dennis Diaz, Juanfang Ruan, Daniel Luque, Bernadette Saunders, Lyndsey E. Collins-Praino, Lars Ittner, Andrew Care

## Abstract

Encapsulins are prokaryotic self-assembling protein nanocages with promise as nanovaccine scaffolds. Their utility as modular platforms require tolerance to surface engineering, high-yield soluble production, formulation stability, and controlled antigen (co-)display. Herein, a previously uncharacterized encapsulin from Alkaliphilus metalliredigens is engineered into a SpyCatcher-decorated nanoscaffold (Am-S) that enables controlled surface display of SpyTagged antigens. Cryo-EM confirms that the native encapsulin forms a *T* = 1 icosahedral nanocage, and that C-terminal SpyCatcher fusion yields Am-S without compromising nanocage assembly, symmetry, or structural integrity. Notably, Am-S exhibits high-yield soluble production in *Escherichia coli*, remains monodisperse after freeze–thaw and extended storage, and supports efficient SpyTagged peptide conjugation for single- and multi-antigen display. As a proof-of-concept, Am-S is functionalized with Alzheimer’s disease-associated amyloid-β and/or hyperphosphorylated tau epitopes to generate single-target nanocages displaying either antigen and dual-target mosaic nanocages co-displaying both. In mice, Am-S antigen display enhances antigen-specific IgG responses relative to free antigens and induces predominantly IgG1-biased humoral immunity. Mosaic nanocages elicit antibodies against both targets, with immune sera selectively recognizing amyloid-β- and phosphorylated tau-associated pathology in ex vivo brain sections from Alzheimer’s disease mouse models. These findings position Am-S as a manufacturable scaffold for developing multi-targeting nanovaccines against complex diseases.

## Introduction

Protein nanoparticles (PNPs), including self-assembling virus-like particles (VLPs) and non-viral protein nanocages, are attractive platforms for nanovaccine development and offer advantages over conventional synthetic nanoparticles. PNPs resemble viruses, presenting repetitive surfaces that can be engineered to display antigens multivalently, enhancing B-cell receptor engagement and inducing adaptive immune responses, including antigen-specific antibodies and T-cell responses. They range from 20 to 200 nm in diameter, supporting lymph node trafficking ^1,2^, and can incorporate intrinsic or engineered T-cell epitopes to further promote humoral and cellular immunity ^3^. They are generally stable, retaining immunogenicity after drying or extended storage ^4^, which could facilitate distribution without strict cold-chain requirements ^5^. PNPs can be produced at scale with favorable safety profiles, and several vaccines based on these platforms are clinically approved against infectious diseases, including hepatitis B, human papillomavirus, malaria ^6^, and COVID-19 ^7^.

Encapsulins are an emerging class of PNPs found in many prokaryotes, where they function as intracellular pseudo-organelles. They self-assemble from identical protein subunits into hollow nanocages that adopt one of three icosahedral architectures defined by triangulation number (T): *T* = 1 (18–24 nm, 60-mer), *T* = 3 (∼32 nm, 180-mer), and *T* = 4 (∼43 nm, 240-mer) ^8–11^. Encapsulins have gained attention as adaptable platforms for nanomedical applications due to their capacity for payload encapsulation and delivery ^12^, intrinsic immunostimulatory activity, and favourable in vivo safety profiles ^13,14^. Their immune-stimulating properties have led to the use of encapsulins as scaffolds for immune priming ^15^ and for antibody discovery and development ^16^. They have also been engineered for antigen display and delivery, with encapsulin-based nanovaccines evaluated preclinically for prophylactic vaccination against infectious diseases, including Epstein-Barr virus ^17^, influenza ^18,19^, Lassa virus ^20^, Severe acute respiratory syndrome coronavirus 2 (SARS-CoV-2) ^14,21,22^, Rotavirus ^23^, African Swine Fever virus ^24–26^, and Salmonella ^27^ (**Table S1**).

Despite recent advances in encapsulin nanovaccine development, several limitations remain incompletely addressed. For instance, there is a reliance on nanocages derived from the bacteria *Thermotoga maritima* (Tm-Enc, *T* = 1) and *Myxococcus xanthus* (Mx-Enc, *T* = 3), restricting access to alternative encapsulins that may offer improved engineerability, immunogenicity, stability, and manufacturability. Moreover, antigen display is often achieved through direct genetic fusion to encapsulin subunits, in which antigen-encapsulin fusion proteins assemble into antigen-decorated nanocages ^14,17–19,23,27–29^. However, the direct fusion of antigens can reduce soluble expression, impair nanocage self-assembly, and decrease production yields, thereby constraining scalable manufacture. To overcome such limitations, split-protein coupling systems (e.g., SpyTag/SpyCatcher ^15,20–22,24–26^ or split inteins ^16^), have been used as modular “plug-and-play” approaches for post-assembly covalent conjugation of antigens to the exterior surface of encapsulins, enabling the generation of nanovaccine candidates. Nevertheless, encapsulin-based nanovaccine development has focused predominantly on monovalent display of foreign antigens for prophylactic vaccination against infectious diseases, which induce neutralising antibody responses against a single non-self target protein (e.g., SARS-CoV-2 spike receptor-binding domain). This narrow application scope contrasts with the broader potential of encapsulins as immunogenic scaffolds. Unlike virus-derived PNPs, encapsulins have not been directly linked to human infection and exhibit minimal sequence similarity to human proteins, reducing the theoretical risks of pre-existing immune tolerance and cross-reactive autoimmunity. These properties suggest that encapsulins could enable prophylactic and therapeutic nanovaccines for complex diseases by targeting antigenic fragments derived from pathological self-proteins, where controlled disruption of B-cell tolerance is required.

Multifactorial diseases provide a relevant context in which multi-targeting encapsulin nanovaccines may be advantageous. Alzheimer’s disease is a complex neurodegenerative disorder characterized by two major pathological protein species, amyloidl7lβ (Aβ) and hyperphosphorylated tau (pTau), which act synergistically to drive disease progression and associated cognitive decline ^30^. Both proteins are established immunotherapeutic targets in passive antibody approaches, including FDA-approved monoclonal antibodies with modest clinical benefit, known safety concerns, and high costs (e.g., Aβ-targeting lecanemab) ^31,32^, as well as in active vaccination strategies ^33,34^. However, most prophylactic and therapeutic vaccine programs have generally targeted a single antigen (Aβ or pTau), and none have prevented cognitive decline. Accordingly, simultaneous co-targeting of Aβ and pTau provides a rational proof-of-concept framework to evaluate multivalent display of endogenous self-antigens on encapsulin-based nanovaccines.

In this article, we present the identification and characterization of a new encapsulin from the bacterium *Alkaliphilus metalliredigens* and its engineering into a SpyCatcher-decorated nanoscaffold for modular co-display of SpyTagged antigens. The nanoscaffold was evaluated for manufacturing-relevant production, stability, and its capacity to elicit antigen-specific antibody responses. As a proof-of-concept, we used the encapsulin nanoscaffold to compare mono- and multivalent display formats of Aβ- and pTau-derived self-antigen peptides and assessed antibody recognition and dual-targeting of human Aβ and pTau in brain tissue from Alzheimer’s mouse models.

## Results and discussion

### Identification and characterization of a novel encapsulin derived from the bacterium *Alkaliphilus metalliredigens*

Bioinformatic surveys have identified over 6,000 putative encapsulin-like systems within bacterial and archaeal genomes ^35^. Among these, the extremophilic firmicute *Alkaliphilus metalliredigens* encodes a predicted *T* = 1 encapsulin system (AmEnc) (**Table S2**). Isolated from borax-contaminated leachate ponds, *A. metalliredigens* thrives in alkaline, hypersaline environments containing high concentrations of heavy metals ^36^. We hypothesized that an encapsulin nanocage derived from such an extremophile would exhibit enhanced biophysical and physicochemical stability, properties highly desirable for the development of new robust biologics e.g., nanovaccines.

Other putative proteins from *A. metalliredigens* have been successfully produced for characterization using *Escherichia coli* as a recombinant host ^36^. Accordingly, we expressed the gene encoding the AmEnc nanocage in *E. coli* (**Fig. 1a**), purified it via sequential size-exclusion (SEC) and anion-exchange chromatography (**Fig. S1**), and subjected it to biophysical characterization (**Fig. 1**). SDS-PAGE analysis revealed a single band near the theoretical size of the AmEnc subunit (29.9 kDa), confirming successful expression and purification (**Fig. 1b**). Under native-PAGE conditions, AmEnc formed a high-molecular-weight band, consistent with self-assembly into the expected nanocage (**Fig. 1c**). Cryo-electron microscopy (cryo-EM) revealed that AmEnc assembles into uniform isometric hollow nanocages with the expected icosahedral shape (**Fig. 1d**). Three-dimensional refinement yielded a reconstruction at 3.1 Å resolution (**Fig. 1e**) (PDB: 22NB; **Table S3**) which showed an icosahedral shell composed of 60 identical subunits arranged as 12 pentameric vertices with *T* = 1 icosahedral architecture, with an external diameter of 21.2 nm and an internal diameter of 15.6 nm, resulting in a luminal cavity of approximately 760 nm³ (**Fig. 1f**). Comparison across well-characterized encapsulins of varying triangulation classes (*T* = 1, *T* = 3, *T* = 4) indicated that the AmEnc subunit conformation is most similar to the *T* = 1 TmEnc subunit (PDB: 3DKT), adopting the characteristic HK97-like fold comprising A- and P-domains and an E-loop (**Fig. S2a**), with a lumen-facing N-terminus and an exterior-facing C-terminus (**Fig. 1g**). The shell contains multiple pores located at the symmetry-axis typical of encapsulins, including a five-fold pore (∼5 Å diameter) at the pentameric axis (**Fig. 1h**), as well as pores at the 2l7lfold and 3l7lfold axes, and a 2l7lfold adjacent pore (**Fig. S3**). All pores display predominantly negative to neutral electrostatic potentials on their luminal and exterior surfaces; this is consistent with their role in mediating the selective flux of small molecules and ions across the encapsulin shell (**Fig. 1h & Fig. S3**).

**Figure 1.**
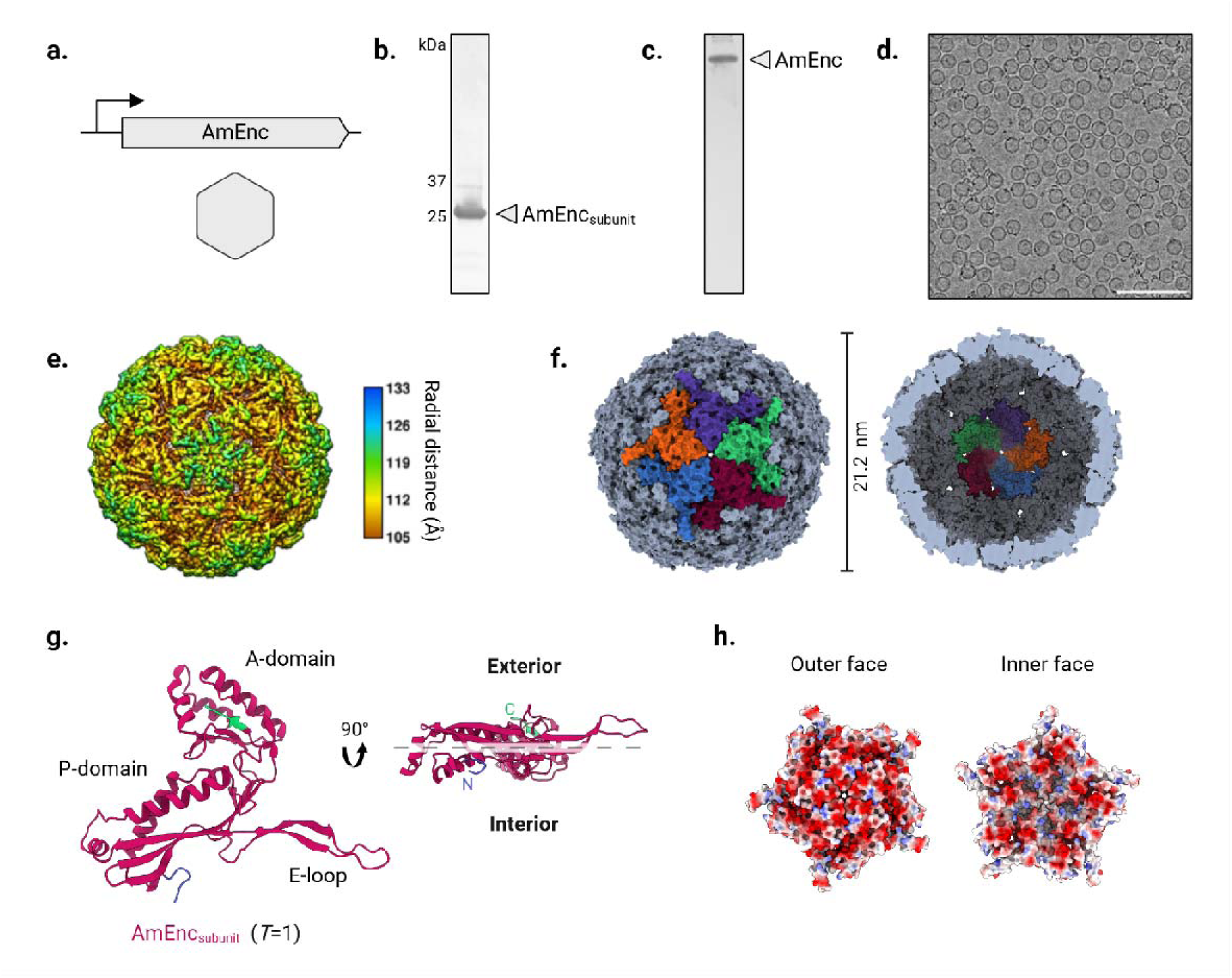
Biophysical characterization and cryo-EM structure of the AmEnc nanocage. (**a**) Schematic of the AmEnc expression construct. (**b**) SDS-PAGE analysis of purified AmEnc showing the expected 29.9 kDa subunit. (**c**) Native-PAGE confirming self-assembly into high-molecular-weight nanocage complexes. (**d**) Representative cryo-EM micrograph of self-assembled icosahedral AmEnc shells (scale bar = 100 nm). (**e**) Cryo-EM reconstruction of assembled AmEnc at 3.1 Å resolution, shown as a surface representation and colored by radial distance (external view). (**f**) Surface-shaded representation of the outer (**left**) and inner (**right**) surfaces of the AmEnc density map revealing a hollow 60-subunit *T* = 1 icosahedral shell with external and internal diameters of 21.2 nm and 15.6 nm, respectively. (**g**) Atomic model of the AmEnc subunit highlighting the canonical HK97-like fold, including the A-domain, P-domain, and E-loop, with the N-terminus (blue) facing the lumen and the C-terminus (green) facing the exterior. (**h**) Electrostatic potential surface of the five-fold symmetry pore viewed from the exterior (**left**) and interior (**right**), with negative and positive potentials shown in red and blue, respectively.

Functionally, AmEnc is predicted to be a classical Family 1 encapsulin, many of which are characterized by the encapsulation of ferritin-like proteins (FLPs) to manage iron homeostasis ^35^ (**Fig. S4**). In support of this, purified AmEnc nanocage exhibited a distinct yellow coloration which, alongside a conserved Trp90 residue, suggests the nanocage surface binds flavin cofactors (**Fig. S4b**). These cofactors are often utilized by Family 1 systems to facilitate the redox chemistry required for efficient iron mineralization. Furthermore, the AmEnc operon encodes a downstream FLP with a C-terminal encapsulation signal (ESig) that mediates targeted encapsulin packaging ^37^. Together, PAGE analysis and cryo-EM confirmed that AmEnc encapsulates multimeric FLPs (**Fig S4c–f**), while TEM imaging following exposure to aqueous ferrous iron revealed the formation of electron-dense mineral cores (**Fig. S4g**). Taken together, these findings experimentally confirm the structure and functionality of the AmEnc system as an iron-sequestering classical Family 1 encapsulin.

### Integration of the SpyTag/SpyCatcher coupling system into AmEnc

Having established the structure and assembly of the newly characterized AmEnc nanocage, we next sought to develop it into a modular “plug-and-play” nanoscaffold for antigen display using the SpyTag/SpyCatcher coupling system ^38–41^. SpyTag (ST) and SpyCatcher (SC) have each been incorporated into different encapsulin architectures to mediate molecular surface display of various antigens (**Table S1**).

Incorporation of the SC domain (12.3 kDa) onto the encapsulin surface is often preferred as it results in a more versatile nanovaccine scaffold. SC decoration mediates the coupling of antigens tagged with the small ST peptide (1.1 kDa), which can be generated via rapid chemical synthesis. This is particularly advantageous for antigens with post-translational modifications (PTMs) (e.g., glycosylation, phosphorylation) that can be introduced site-specifically during synthesis or via recombinant expression in a eukaryotic host. Furthermore, the small size and flexible structure of ST-tagged peptide antigens may reduce steric interference, maximizing coupling efficiency and antigen density, while also enabling the co-display of different antigens to create multivalent nanovaccines for multi-targeted intervention against complex diseases. However, despite these advantages, incorporating the larger SC into encapsulin architectures could be expected to present challenges akin to direct genetic fusion (i.e. impaired folding of the nanocage subunits, leading to protein insolubility, inclusion body formation, and reduced yields).

To construct and evaluate the feasibility of a SpyCatcher-decorated AmEnc (Am-S) nanoscaffold, we genetically fused the SC sequence to the surface-exposed C-terminus of the AmEnc subunit via a flexible linker containing a polyhistidine tag (His-tag) (**Fig. 2a**). This configuration facilitates His-tag–based purification while ensuring that the SC domain is sufficiently projected from the nanocage surface. The design was intended to minimize disruption to encapsulin self-assembly while maximizing accessibility for subsequent antigen coupling.

**Figure 2.**
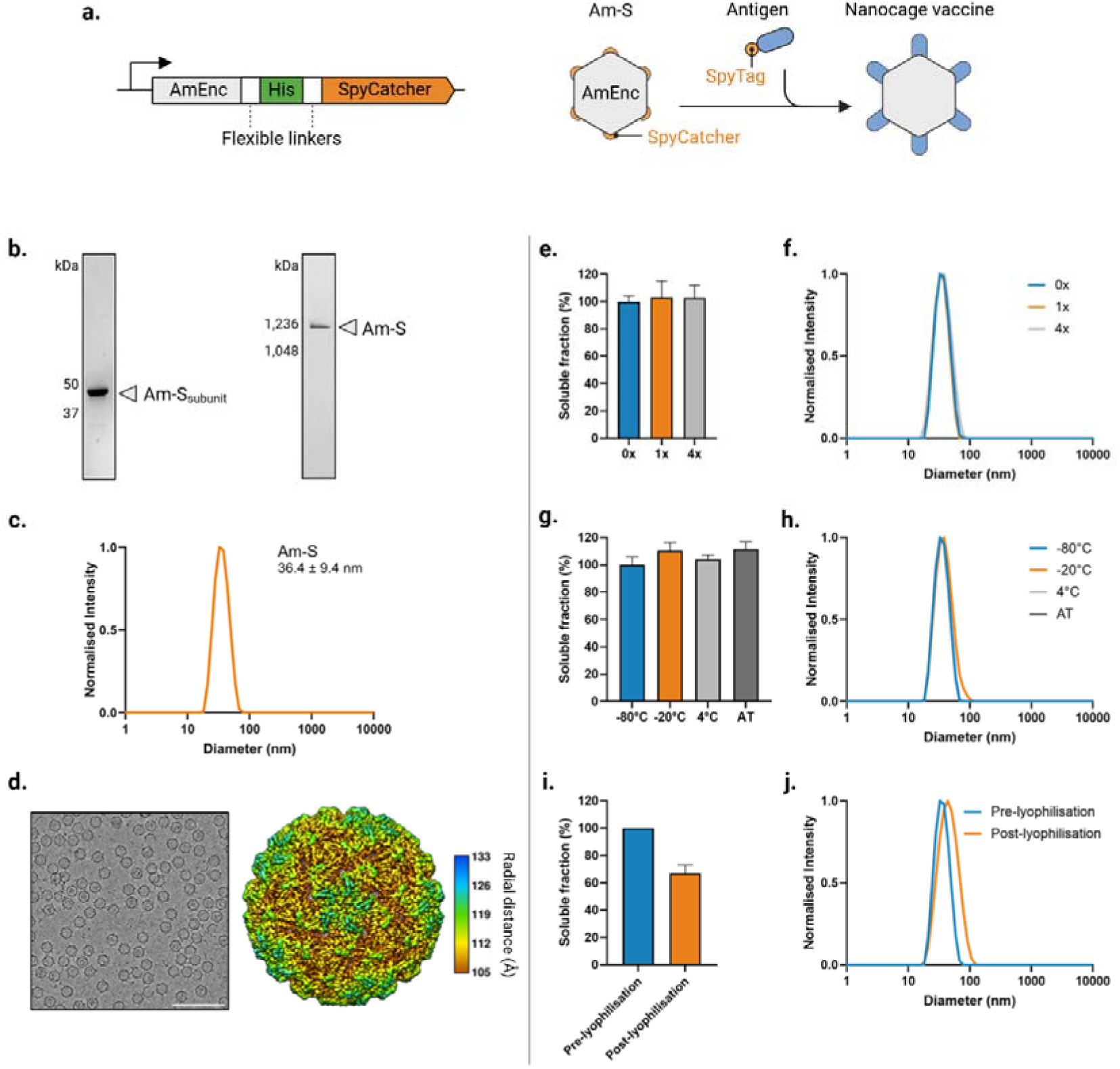
Design, production, biophysical characterization, and storage stability of the SpyCatcher-decorated AmEnc nanocage (Am-S). (**a**) Schematics depicting (**left**) the Am-S genetic construct encoding the AmEnc subunit (grey) fused at the C-terminus to SpyCatcher (orange) via a linker comprising a His-tag (green) flanked by flexible (GGGS)*_n_* spacers (white); and (**right**) covalent coupling of SpyTagged antigens to the surface of the self-assembled Am-S scaffold, generating an antigen-displaying nanovaccine. (**b**) PAGE analysis of Am-S purified by sequential IMAC and SEC: (**left**) SDS-PAGE showing the Am-S subunit (∼44 kDa); (**right**) native PAGE verifying nanocage assembly. (**c**) DLS analysis demonstrating monodisperse Am-S nanocages with a mean hydrodynamic diameter of 36.4 ± 9.4 nm. (**d**) (**left**) Cryo-EM micrograph showing Am-S self-assembly into nanocage structures (scale bar = 100 nm) (**right**) 3D reconstruction at 2.58 Å resolution (external view) confirming high-fidelity assembly into 21.2 nm particles with *T* = 1 icosahedral symmetry. (**e–j**) Storage stability of Am-S. (**e**) Solubility after 1 and 4 freeze-thaw cycles; untreated material (0) was defined as 100% soluble. (**f**) DLS analysis of samples in (e). (**g**) Solubility after storage at the indicated temperatures for 6 weeks; samples stored at −80 °C were defined as 100% soluble. (**h**) DLS analysis of samples in (g). (**i**) Solubility before and after lyophilisation and storage at ambient temperature for 1 day; pre-lyophilisation material was defined as 100% soluble. (**j**) DLS analysis of samples in (i). Soluble Am-S fractions were isolated by centrifugation and quantified by SDS-PAGE densitometry (mean ± SD, *n* = 3).

Am-S was recombinantly expressed in *E. coli*, yielding substantial amounts of soluble protein. Notably, under identical expression conditions, an equivalent construct encoding a SpyCatcher-decorated *T. maritima* encapsulin (Tm-S) exhibited near-complete insolubility, with only trace amounts of soluble protein detected (**Fig. S5**). Am-S showed markedly higher soluble protein yield that Tm-S, indicating that the AmEnc scaffold is more tolerant of C-terminal fusion and surface display of large protein domains that its more commonly used TmEnc counterpart.

Soluble Am-S was purified via immobilised metal affinity chromatography (IMAC) with integrated on-column endotoxin removal ^42^, followed by a SEC polishing step to isolate self-assembled nanocages from free subunits (**Fig. S6a & b**). Following purification, host-cell nucleic acid contamination was minimal (A_260nm_/A_280nm_ = 0.6) ^43^. SDS-PAGE confirmed the successful purification of the Am-S subunit, revealing a distinct band consistent with its theoretical molecular weight of 43.9 kDa, while densitometric analysis determined a purity of >90% (**Fig. S6c & Fig. 2b, left**). Under these conditions, Am-S yielded ∼200 mg/L of purified protein, dramatically exceeding yields obtained for Tm-S (data not shown) and representing a 10-fold higher yield than reported for other modified encapsulins incorporating the SC/ST system ^21,44^.

To assess whether SC decoration affected nanocage self-assembly, we performed detailed biophysical characterization. Native-PAGE analysis of Am-S samples visualised high-molecular-weight species consistent with nanocage assembly (**Fig. 2b, right**), while DLS measurements confirmed nanocages with a mean hydrodynamic diameter of 36.4 ± 9.4 nm and high monodispersity (PDI = 0.05) (**Fig. 2c**). Cryo-EM imaging showed that Am-S forms uniform, hollow nanocages with overall morphology similar to that of unmodified AmEnc (**Fig. 2d, left**). Subsequent 3D refinement yielded a 2.58 Å reconstruction of the encapsulin shell, revealing that the Am-S capsid retains the *T* = 1 icosahedral architecture and subunit fold of AmEnc, assembling as a high-fidelity 60-mer nanocage (EMDB: 80547; **Fig. 2d, right, Table S3**). The SC domains were not resolved as discrete ordered features, but appeared as diffuse peripheral density on the exterior surface of the shell (**Fig. S7**), consistent with linker-mediated flexibility. These results demonstrate that C-terminal SC incorporation does not measurably perturb AmEnc capsid assembly, structural integrity, or symmetry, while generating a surface-functionalized nanoscaffold suitable for subsequent coupling to SpyTagged antigens.

Vaccines and other biopharmaceuticals typically require cold storage and rely on cold-chain supply chains, which are costly, limit widespread distribution, and are susceptible to breaches that can render products ineffective or unsafe ^5^. Accordingly, encapsulin-based nanovaccines that tolerate repeated freeze-thaw cycles and storage at broad temperature ranges would offer logistical and commercial advantages.

We have previously established that encapsulins have high thermal stability and resilience to extreme pH, chemical denaturants, and multiple freeze-thaw cycles ^13^. In similar reports, SpyTag-decorated MxEnc and TmEnc nanoscaffolds have exhibited prolonged stability at ambient temperature, maintaining nanocage structure for up to 12 months and 150 days, respectively ^24,44^. However, it should be noted that the MxEnc-SpyTag nanoscaffolds showed reduced colloidal stability upon heat treatment, with reported PDI values of 0.32 at 40 °C and 0.56 above 40 °C ^44^. Khaleeq et al. further demonstrated that an MxEnc-SpyTag nanovaccine displaying receptor-binding domain (RBD) antigens was stable after storage for up to 4 weeks at 4 °C and 2 weeks at 37 °C ^21^, without loss of antigenicity. Despite these insights, encapsulin stability under other storage conditions relevant to vaccine development, such as lyophilisation, remains unexplored. Therefore, to assess the potential utility of the Am-S scaffold in nanovaccine applications, we evaluated its stability under a range of storage conditions in phosphate-buffered saline (PBS) without added stabilizing excipients (**Fig. 2e–j**).

After multiple freeze-thaw cycles at -80 °C, SDS-PAGE analysis revealed that Am-S retained its solubility (**Fig. 2f**) without any observable degradation (**Fig. S8a**), while DLS measurements confirmed no apparent change in its hydrodynamic diameter nor any aggregate formation (**Fig. 2g**). This is consistent with our previous results showing TmEnc as resilient to multiple freeze-thaw cycles ^13^. To test longer-term storage, Am-S was stored at -80 °C, -20 °C, 4 °C, and ambient temperature (20–25 °C) for 6 weeks and retained both solubility (**Fig. 2h**) and structure without aggregation (**Fig. 2i**) at all temperatures. No degradation was observed after storage at -80 °C, -20 °C, or 4 °C, however, after storage at ambient temperature, a faint band corresponding to the size of AmEnc without SpyCatcher (∼30 kDa) appeared (**Fig. S8b**), suggesting a small degree of proteolytic activity. This is likely from trace contamination, which could be remedied with further purification optimisation. Faint higher molecular weight bands were also observed after storage at 4 °C and ambient temperature and likely represent Am-S multimers that did not completely denature during SDS-PAGE sample preparation (**Fig. S8b**). Intact multimers in SDS-PAGE have been similarly reported in variants of the highly thermostable TmEnc ^45,46^. Given that the thermal stability of AmEnc or Am-S at high temperatures was not evaluated in this study, further investigation is required to understand why certain storage conditions may make Am-S more resilient to denaturation.

Lyophilization can improve the long-term stability of protein-based vaccine formulations; however, the process introduces cryogenic and dehydrative stresses that can induce structural perturbation, degradation, and/or aggregation, potentially compromising vaccine immunogenicity ^47^. To assess the effects of rapid freeze-drying, Am-S underwent lyophilisation prior to storage at ambient temperature. Upon reconstitution the following day, 67% of soluble protein remained (**Fig. 2j**), with only minimal aggregation or swelling, as indicated by an increase in hydrodynamic diameter (**Fig. 2k**), and no signs of degradation observed (**Fig. S8c**). These results indicate that Am-S is a highly robust nanoscaffold for antigen display, resilient to aggregation and maintains solubility after multiple freeze-thaw cycles and prolonged storage at the tested temperatures. Nevertheless, lyophilisation induced a moderate material loss and detectable aggregation and/or swelling. Future work could therefore focus on optimising freezing and drying parameters and evaluating the inclusion of lyoprotectants, such as trehalose or sucrose, to further enhance Am-S stability ^47^.

The above work shows that SpyCatcher integration does not compromise the self-assembly or symmetry of the AmEnc nanocage. Am-S exhibits high solubility and production yields that exceed those reported for other encapsulin-based display systems. Moreover, the scaffold remains monodisperse and structurally stable following repeated freeze–thaw cycles, extended storage, and lyophilisation. Collectively, these properties support Am-S as a robust scaffold for the potential scalable construction and manufacture of nanovaccines.

### Am-S serves as a nanoscaffold for antigen (co-)display

To our knowledge, encapsulin-based nanovaccine platforms have thus far been mostly limited to monovalent antigen display and the targeting of single prophylactic or therapeutic targets (**Table S1**). Recently, TmEnc was genetically modified for multivalent display of *Salmonella* surface membrane epitopes ^27^. While this is initially encouraging, one epitope, displayed only in the monovalent configuration, was assessed in vivo, while the impact of multiple different antigens on immune response remains to be determined. We therefore investigated whether the Am-S nanoscaffold could simultaneously conjugate and co-display multiple SpyTag-tagged peptide antigens, including those bearing complex PTMs.

As a proof-of-concept, we focused on Alzheimer’s disease (AD), a heterogeneous neurodegenerative disorder defined by two major pathological protein species: intracellular hyperphosphorylated tau (pTau) and extracellular amyloid-β (Aβ). Recent evidence indicates a pathological synergy between pTau and Aβ, in which they exacerbate each other’s toxicity to drive AD progression ^30^. Accordingly, we selected well-characterized antigens from each protein that have shown promise in both passive and active immunotherapy approaches. These included (i) the pS396/pS404 pTau epitope, the targeting of which via active immunotherapy has been shown to reduce tau pathology ^48,49^, tauopathy-related motor impairments ^50,51^, and cognitive impairment ^52^ in preclinical studies, and has now entered clinical trials ^51^; and (ii) the Aβ_1-6_ sequence, a highly immunogenic region of the Aβ N-terminus that is targeted by FDA-approved passive anti-Aβ monoclonal antibodies ^53^.

To enable coupling to Am-S, we synthesized two peptides with N-terminal SpyTags: a tau_389-408_ peptide bearing site-specific phosphorylation at Ser396 and Ser404 (S-pTau); and an Aβ_1-6_ peptide (S-Aβ) (**Fig. 3a**). Importantly, the use of these antigens not only allows the adaptability of the Am-S nanoscaffold to be assessed, but also helps further the development of multi-targeting AD vaccines.

**Figure 3.**
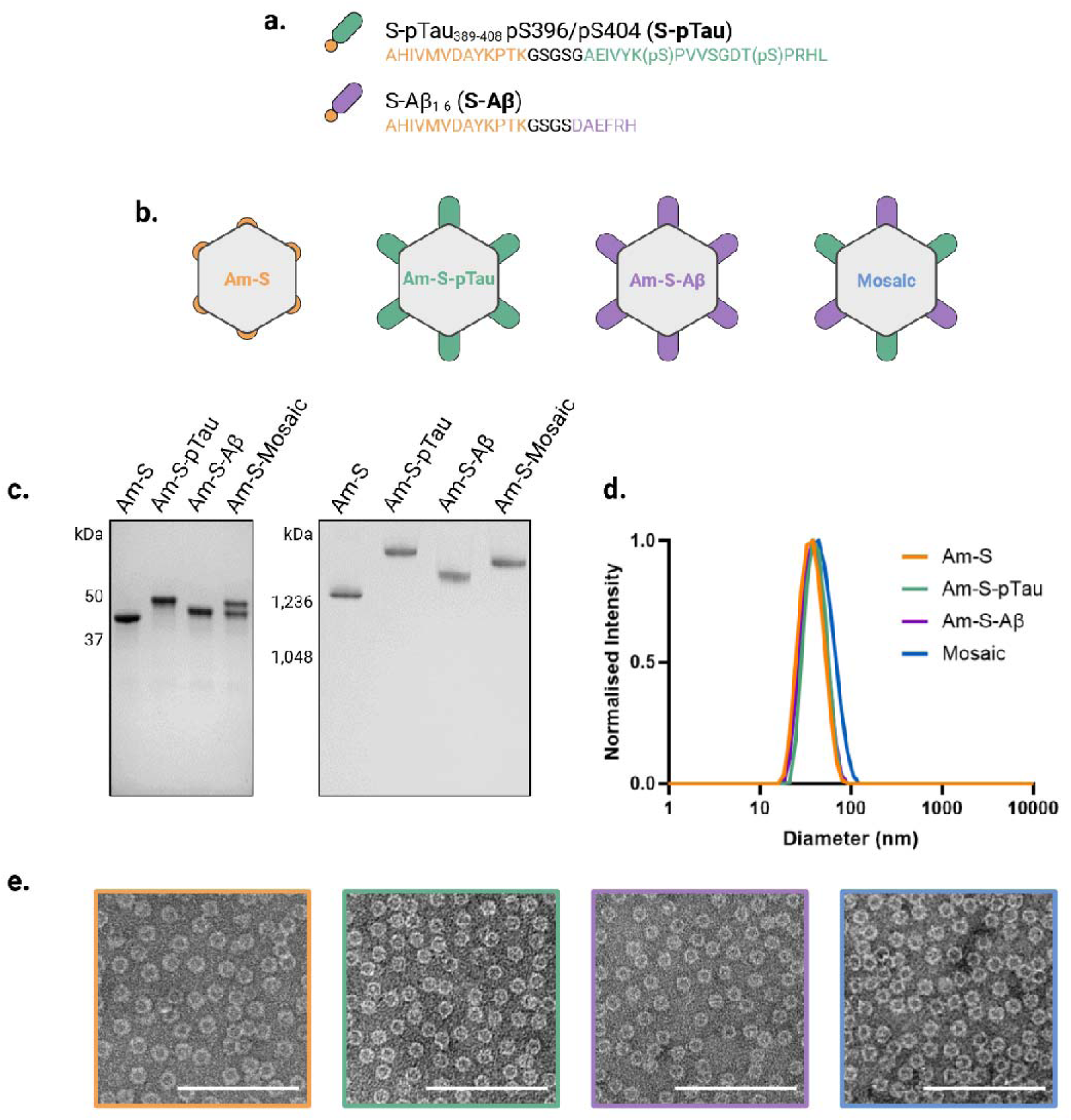
(Co-)conjugation of SpyTagged AD antigens onto the Am-S nanoscaffold. (**a**) Schematic of peptide antigens with an N-terminal SpyTag (orange) linked via a flexible (GS)n spacer (black) to peptide antigens derived from pTau (S-pTau; green) or Aβ (S-Aβ; purple). (**b**) Conceptual illustration of unconjugated and conjugated Am-S, including the bare nanoscaffold (Am-S), monovalent nanocage formats bearing S-pTau (Am-S-pTau) or S-Aβ (Am-S-Aβ), and a multivalent “mosaic” nanocage bearing both antigens. (**c**) PAGE assessment of SpyTag/SpyCatcher-mediated conjugation, with (**left**) SDS-PAGE showing covalent coupling of antigen(s) to the Am-S subunit, and (**right**) non-denaturing native PAGE indicating antigen (co-)display on the assembled Am-S nanocage. (**d**) DLS characterisation of hydrodynamic diameter and dispersity of the (co-)conjugated nanocages. (**e**) Negatively stained TEM images of Am-S (orange), Am-S-pTau (green), Am-S-Aβ (purple), and mosaic (blue) nanocage formats; Scale bars = 200 nm.

The Am-S nanocage consists of 60 identical SpyCatcher-bearing subunits, providing up to 60 potential conjugation sites per cage for controlled antigen attachment. As illustrated in **Figure 3b**, we sought to exploit this modularity to construct both monovalent nanocages displaying either S-pTau (Am-S–pTau) or S-Aβ (Am-S–Aβ), as well as multivalent “mosaic” nanocages co-displaying both antigens. Optimal conjugation conditions were initially identified by screening a range of molar ratios of Am-S to Spy-tagged antigens, with reactions performed for 16 h at 4 °C (data not shown).

For monovalent Am-S–pTau nanocage construction, Am-S cages were reacted with excess S-pTau at a 1:4 molar ratio. Bioconjugation of the Am-S subunit with S-pTau was visualized by SDS-PAGE, which revealed a distinct band shift from the unmodified Am-S subunit (43.9 kDa) to the expected molecular weight of the conjugated Am-S–pTau subunit (47.9 kDa) (**Fig. 3c, left)**. Densitometric analysis confirmed complete conversion of the Am-S subunit, indicating high conjugation efficiency and saturation of all available SpyCatcher sites on the nanocage surface. Intact mass increases relative to bare Am-S nanocages were also observed via blue native PAGE (**Fig. 3c, right**), together with a modest increase in hydrodynamic diameter measured by DLS (**Fig. 3d**), indicative of surface decoration with S-pTau antigens. TEM imaging showed that Am-S-pTau nanocages retained their assembled structure and morphology, with no evidence of aggregation (**Fig. 3e**). Comparable results were obtained for monovalent Am-S-Aβ nanocages, generated by reacting Am-S with excess S-Aβ at a 1:3 molar ratio (**Fig. 3c–e**).

To construct mosaic nanocages, we simultaneously mixed Am-S, S-pTau, and S-Aβ at a 1:4:1 molar ratio. SDS-PAGE revealed two distinct bands corresponding to Am-S subunits conjugated to the differently sized antigens to S-pTau (47.9 kDa) and S-Aβ (46.4 kDa) (**Fig. 3c, left**). Densitometric analysis determined high conjugation efficiency with approximately equivalent proportions of Am-S subunits conjugated to each antigen using the 1:4:1 ratio (**Fig. S9**), confirming successful co-display on mosaic nanocages. Blue native PAGE showed a singular band with a molecular weight between that of Am-S-pTau and Am-S-Aβ nanocages (**Fig. 3c, right**), consistent with the formation of a homogeneous population of mosaic nanocages arising from equimolar co-display with two antigens of differing molecular weights. As expected, DLS showed a modest increase in hydrodynamic diameter without aggregation (**Fig. 3d**), and TEM confirmed preserved nanocage morphology (**Fig. 3e**).

Overall, the Am-S nanoscaffold provided an efficient and controllable platform for the high-density (co-)display of S-pTau and S-Aβ antigens. These antigens were coupled either individually or simultaneously without detectable aggregation, demonstrating the utility of Am-S as a modular antigen display platform. Precise positioning and spacing of antigens on the scaffold surface can enhance the immunogenicity of nanovaccines by promoting efficient B-cell receptor (BCR) crosslinking ^1,2^. Moreover, high antigen density is a critical factor in stimulating class-switching from early-stage antigen-specific immunoglobulin M (IgM) to IgG isotypes, which is associated with longer-lasting immunity and has been observed with other antigen-displaying protein cages ^54^. Dense antigen coverage can also help circumvent carrier-induced epitopic suppression, a phenomenon in which the scaffold itself elicits an immune response that competes with the displayed antigens. By effectively masking the scaffold surface, high-density antigen display may prevent carrier-specific antibodies from interfering with the induction of the desired anti-antigen response ^55^.

### Am-S nanoscaffold enhances antigen-specific immune responses

To serve as an effective scaffold in nanovaccines, Am-S must exhibit intrinsic immunostimulatory properties that amplify immune responses specific to antigens displayed on its surface. Several encapsulin nanocages demonstrate inherent immunogenicity and have been successfully applied to safely boost antigen-specific immunity in nanovaccine formulations (**Table S1**). We therefore proceeded to characterise the immunostimulatory properties of the newly developed Am-S scaffold.

Using the monovalent Am-S-pTau as a model nanovaccine, we conducted a preliminary screen of the immune and safety profiles of the Am-S platform (**Fig. 4a–c**). C57BL/6J mice (*n* = 4 per group) were subcutaneously injected with either Am-S-pTau, equivalent doses of free S-pTau antigen, Am-S alone, or adjuvant control. Primary injections were supplemented with complete Freund’s adjuvant (FA), followed by booster injections with incomplete FA 14-days later (**Fig. 4a**). FA has previously been used as an adjuvant in other preclinical studies with encapsulins ^18,24–26^. An additional group received Am-S-pTau without FA to evaluate scaffold immunogenicity in the absence of extrinsic adjuvant. As shown in **Fig. 4b**, terminal sera analysis via ELISA revealed that Am-S-pTau/FA elicited significantly higher total anti-pTau IgG levels than all other controls. In contrast, S-pTau with FA or non-adjuvanted Am-S-pTau induced minimal antibody responses, indicating that S-pTau alone is poorly immunogenic. These initial results show that conjugation and display of S-pTau on Am-S, supplemented with FA, synergistically enhances antigen-specific immune responses.

**Figure 4.**
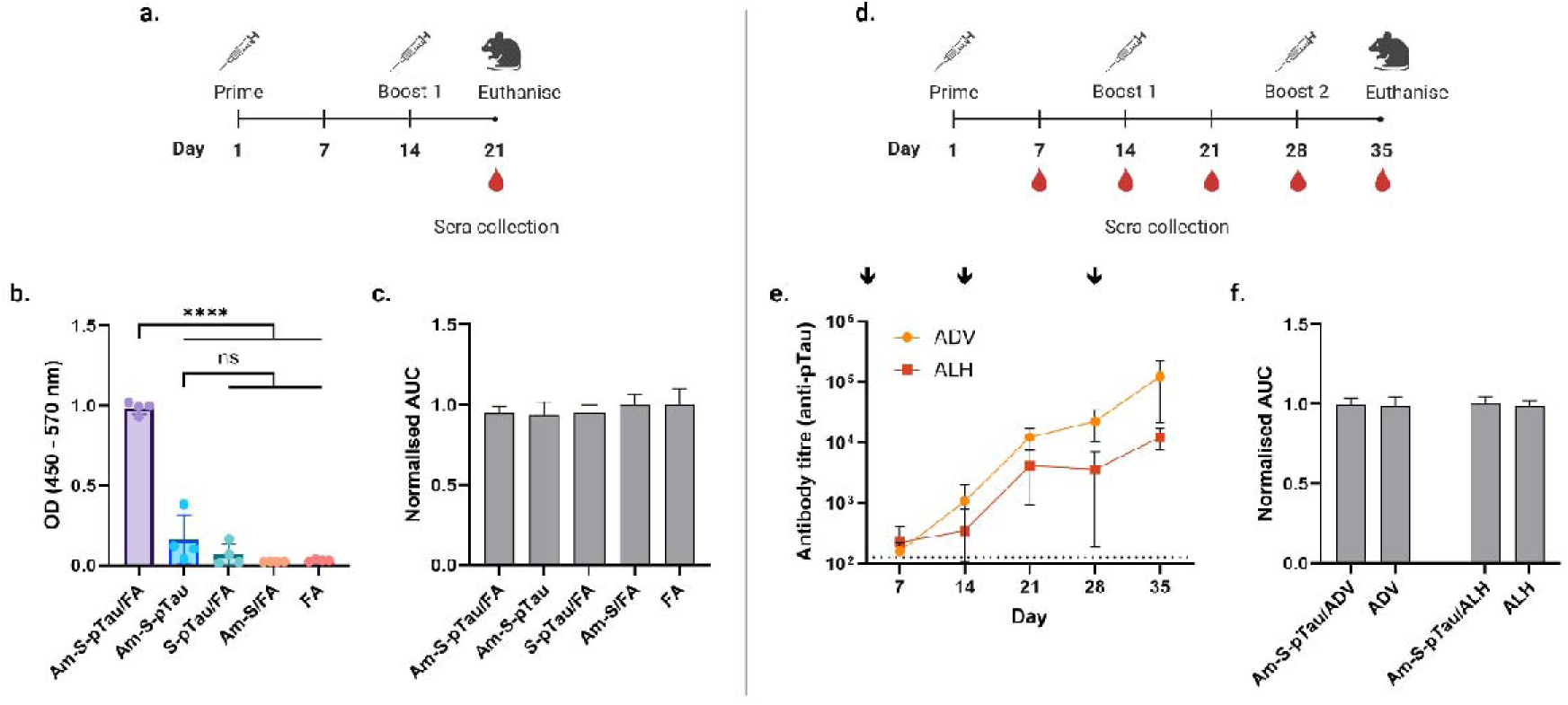
Immunogenicity and tolerability of monovalent Am-S-pTau nanovaccine supplemented with different adjuvants. (Left panel) Freund’s adjuvant (FA). (**a**) Immunisation schedule for C57BL/6J mice (*n* = 4 per group) with end-point sera collection. (**b**) ELISA measurement of anti-pTau total IgG levels in terminal sera from mice receiving Am-S-pTau and controls. (**c**) Body-weight change of mice over time-course in (b), expressed as normalised area under the curve (AUC). (**Right panel**) AddaVax™ and Alhydrogel®. (**d**) Immunisation and sera collection schedule for mice receiving Am-S-pTau formulated with ADV or ALH (*n* = 4 per group). (**e**) Anti-pTau IgG titres in sera over time. Black arrows indicate immunization days. (**f**) Body-weight change over time in (e). Data are mean ± SD. ****P < 0.0001; ns, non-significant (> 0.05). Statistical analyses: one-way ANOVA with Tukey’s multiple comparisons for (b,c,f); two-way mixed-effects ANOVA with time and adjuvant as factors for (e).

Immunisation (with any formulation) did not cause weight loss (**Fig. 4c**) or observable behavioural changes (e.g., hunched posture, decreased activity), nor did it induce skin irritation or signs of acute inflammatory shock or severe hypersensitivity, nor systemic inflammation (**Fig. S10a & b**) indicating a favourable safety profile for Am-S. These findings are consistent with previous in vivo studies of encapsulin nanocages ^13^. However, it is important to note that FA is not approved for human use, and complete FA is a Th1-polarising adjuvant that induces a proinflammatory immune response. In the context of Alzheimer’s vaccine development, inflammatory conditions have been associated with adverse autoimmune events, as observed in the AN-1792 Aβ clinical trial ^56,57^ and in preclinical tau vaccine studies ^58,59^. Consequently, next-generation AD vaccines typically employ adjuvants that elicit a less inflammatory, Th2-type antigen-specific response to minimise the risk of harmful T cell autoreactivity.

With this in mind, the immunogenicity of Am-S-pTau was next evaluated in a comparative experiment using the less inflammatory and more clinically relevant adjuvants AddaVax^TM^ (ADV) and Alhydrogel^®^ (ALH) (**Fig. 4d–f**). ALH is an analogue of the Th2-biased adjuvant Alum, the most widely used adjuvant for human vaccines ^60^; whereas ADV is a squalene-based oil-in-water nanoemulsion analogous to the MF59 adjuvant licensed for human use, which predominantly elicits a Th2 immune response with a weaker Th1 component ^61^. Sepivac SWE^TM^ is identical to ADV and has been used as an adjuvant in previous Enc vaccine studies ^19^ ^21^, as has ALH ^23^.

C57BL/6J mice (*n* = 4 per group) were subcutaneously injected with Am-S-pTau nanocages formulated with either ADV or ALH. Mice received a primary injection followed by boosters on days 14 and 28. To monitor immune responses over time, weekly sera were collected via submandibular bleed, followed by a terminal bleed on day 35 (**Fig. 4d**). As per **Fig. 4e**, ELISA analysis of the sera revealed that anti-pTau IgG antibody titres following immunisation with Am-S-pTau adjuvanted with ADV or ALH increased significantly over time (*P = 0.0465). Moreover, Am-S-pTau/ADV elicited higher antibody titres than Am-S-pTau/ALH, showing a strong trend over time approaching statistical significance (P = 0.0520). As expected, neither immunisation nor adjuvant-only controls resulted in weight loss (**Fig. 4f**) or any signs of behavioural or inflammatory toxicity (**Fig. S10 c & d**). Given that Am-S-pTau formulated with ADV induced higher anti-pTau antibody responses than ALH, ADV was selected for subsequent immunisation experiments.

### Am-S nanoscaffold enables single- and dual-targeting nanovaccine development

We next evaluated whether the Am-S platform could support multi-targeting vaccination strategies by formulating a set of single- or dual-targeting nanovaccines for comparative immunogenicity assessment (**Fig. 5a**). Single-targeting formulations employed monovalent nanocages displaying either pTau (Am-S-pTau) or Aβ (Am-S-Aβ) antigens. Dual-targeting formulations comprised either multivalent “mosaic” nanocages co-displaying both pTau/Aβ antigens on a single particle; or a “cocktail” mixture of monovalent Am-S–pTau and Am-S–Aβ nanocages co-administered in equal amounts.

**Figure 5.**
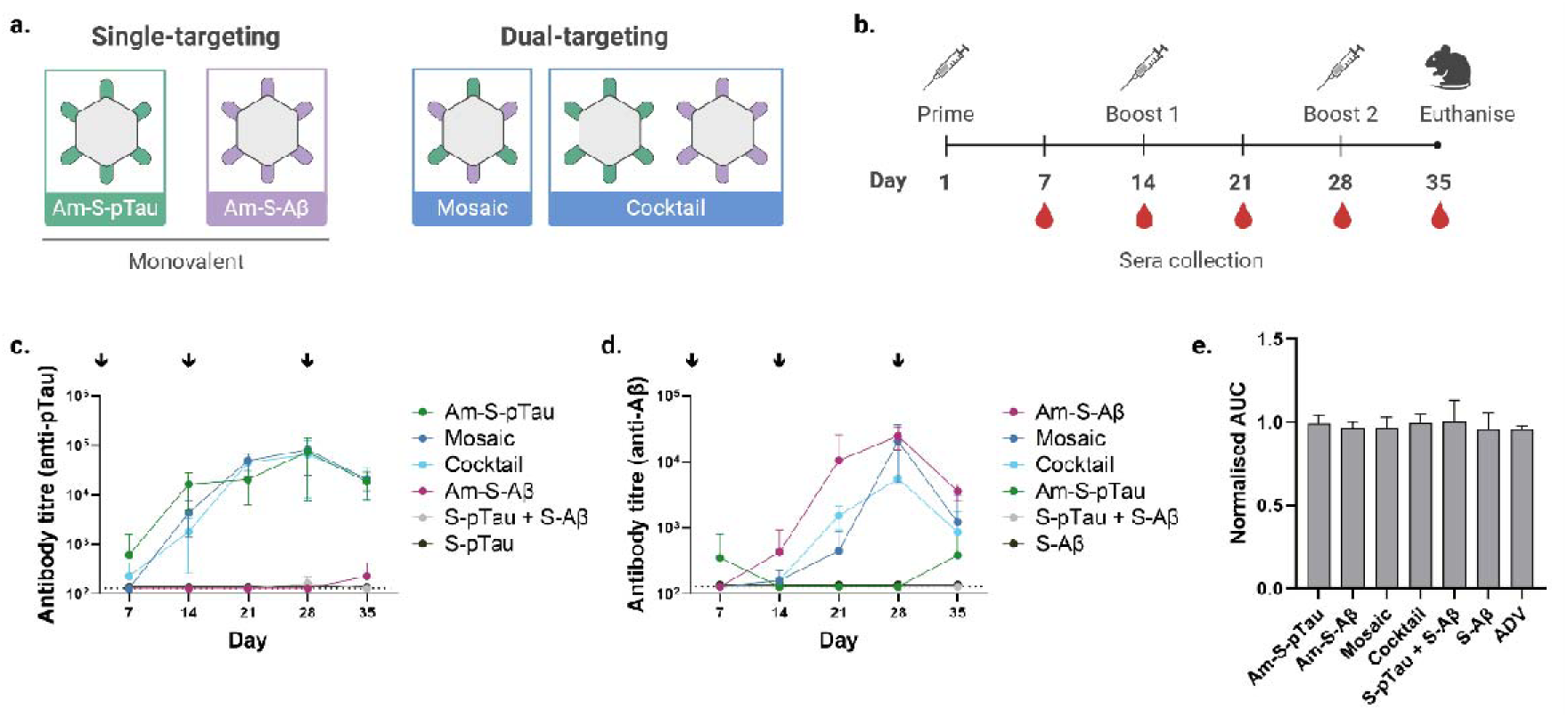
Immunogenicity and tolerability of single- and dual-targeting nanovaccine formulations. (**a**) Schematic of the nanovaccine formats used for immunisation: single-targeting monovalent nanocages, and the dual-targeting multivalent mosaic and cocktail formulations. (**b**) Time-course of immunisation schedule and sera collection for C57BL/6J mice (*n* = 4). (**c**) ELISA measurement of anti-pTau IgG titres in sera over time. (**d**) ELISA measurement of anti-Aβ IgG titres in sera over time. (**e**) Body-weight change of mice in each group over time shown as normalised AUC. Data are mean ± SD. Dotted lines represent the lowest dilution tested. Statistical analyses: two-way ANOVA with repeated measures and Tukey’s multiple comparisons for (c,d), summary statistics are provided in Tables S4 and S5. One-way ANOVA with Tukey’s multiple comparisons for (e). (c,d) Black arrows indicate immunization days.

Herein, C57BL/6J mice (*n* = 4 per group) were immunised subcutaneously with either a single-targeting monovalent nanocage, a dual-targeting mosaic nanocage, or the cocktail formulation (**Fig. 5b**). Control groups received either free antigens administered individually (S-pTau or S-Aβ) or in combination (S-pTau + S-Aβ), or adjuvant alone (ADV). Total pTau and/or Aβ antigen doses were normalised across relevant groups. Mice received three injections at two-week intervals, and sera were collected weekly by submandibular bleed to monitor humoral immune responses over time (**Fig. 5c & d**). Following immunisation, all formulations were well tolerated, with no evidence of weight loss (**Fig. 5e**), behavioural abnormalities, or local skin irritation.

Upon completion of the immunisation regimen, sera collected at each timepoint were analysed by antigen-specific ELISA to quantify titres of antibodies generated against pTau or Aβ. As per **Figure 5c**, anti-pTau antibody titres elicited by Am-S-pTau, mosaic, and cocktail nanovaccines increased over time with a similar trend, peaking at day 28 after two injections before subsiding at day 35. At day 21, both the mosaic and cocktail formulations generated significantly higher antibody levels than the Am-S-Aβ and free-antigen controls (p < 0.05), whereas Am-S-pTau produced higher, but not statistically significant, titres relative to controls. By day 28, Am-S-pTau, mosaic, and cocktail nanovaccines all exhibited significantly greater anti-pTau titres than controls (Am-S-pTau and mosaic, p < 0.0001; cocktail, p < 0.001); no statistically significant differences were observed among the three vaccine formats at any time point (**Table S4**). Similarly, anti-Aβ antibody titres following immunisation with Am-S-Aβ, mosaic, and cocktail nanovaccines increased over time and peaked at day 28 after the second injection, before declining slightly at day 35 (**Fig. 5d**). The Am-S-Aβ nanovaccine induced antibodies earlier than the mosaic and cocktail formats, with significantly higher titres at day 21 relative to controls and the mosaic vaccine (p < 0.05) (**Table S5**). At day 28, both Am-S-Aβ and mosaic nanovaccines elicited significantly higher antibody titres than controls (p < 0.0001), whereas the cocktail formulation showed a higher, but non-significant, trend relative to controls. At this timepoint, anti-Aβ titres did not differ significantly between the Am-S-Aβ and mosaic vaccines; however, both were significantly higher than those induced by the cocktail formulation (p < 0.0001 and p < 0.01, respectively) (**Table S5**). Taken together, these results indicate that coupling pTau and/or Aβ antigens to the Am-S nanoscaffold can enhance their immunogenicity and improve the induction of antigen-specific antibody responses.

Notably, pTau immunogenicity appeared unaffected by co-delivery with Aβ, as the dual-targeting mosaic and cocktail formulations generated pTau-specific titres equivalent to those induced by the single-targeting Am-S–pTau nanovaccine. For anti-Aβ responses, the mosaic nanovaccine achieved titres comparable to the single-targeting Am-S-Aβ vaccine, albeit at later time points, whereas the cocktail formulation induced lower overall anti-Aβ titres, despite delivering equivalent total amounts of both pTau and Aβ antigens. One potential explanation is the shorter length of the Aβ peptide (6 residues) relative to the pTau epitope (19 residues). The pTau-displaying Am-S-pTau component may therefore impose B-cell immunodominance, limiting the ability of the Am-S-Aβ component to elicit anti-Aβ responses during co-delivery. Consistent with this interpretation, prior work on multi-targeting AD nanovaccines using a comparable cocktail strategy reformatted Aβ1–6 as a trimer to increase its effective length and valency ^62^. Under those conditions, anti-Aβ titres matched the monovalent Aβ nanoparticle, while anti-pTau responses exceeded the monovalent pTau comparator. Support for a role of antigen size is further provided by a liposomal nanovaccine study in which Aβ- and pTau-derived peptides of similar length were displayed; no differences were observed between cocktail, mosaic, and monovalent formats ^63^. Future work will investigate the effect of increased Aβ peptide length on the efficiency of Am-S-Aβ and dual-targeting vaccines.

Following the third injection, both anti-pTau and anti-Aβ antibody titres declined modestly from the day 28 peak, with lower levels observed by day 35 across all nanovaccine formulations. This contraction may partially reflect carrier-induced epitope suppression, in which scaffold-directed antibodies generated after earlier doses limit subsequent boosting of the intended antigen-specific response ^55^. Although not tested here, heterologous prime-boost strategies that alternate immunologically distinct carriers can mitigate this effect ^14,64^. For example, during development of a COVID nanovaccine, antigen-specific immunity to SARS-CoV-2 RBD was improved when delivery alternated between encapsulin- and bacteriophage-based protein nanoparticles ^14^. Consistent with this principle, we previously showed that distinct encapsulin species are immunologically orthogonal, inducing carrier-specific antibodies with minimal cross-reactivity ^13^. Whether sequential use of Am-S with alternative encapsulin scaffolds can sustain or enhance antigen-specific titres therefore warrants investigation.

To gain further insight into the immune responses induced by the nanovaccines, sera collected at day 28 that correspond with peak antibody levels were subjected to IgG subclass isotyping (**Fig. S11**). The isotyping of antigen-specific IgG subclasses can be used as an indirect indicator of the type of immune response induced, as cytokines released during a Th1 dominant response (pro-inflammatory) are usually associated with the production of murine IgG2c and IgG2b, and Th2 dominant response (non-inflammatory) produce cytokines associated with IgG1 production. As C57BL/6J mice do not express IgG2a, IgG2c was instead measured as an indirect measure of a Th1 dominant response ^65,66^. All nanovaccine formulations tested preferentially induced the IgG1 isotype, indicative of a non-inflammatory Th2 immune response (**Fig. S11**), suggesting that antigen display on the Am-S scaffold does not strongly bias toward pro-inflammatory responses, a feature advantageous for AD vaccine development.

### Antibodies generated by single- and dual-targeting nanovaccines bind human pathological pTau and/or A**β**

Immunisation with single- or dual-targeting nanovaccine formulations elicited antibodies against pTau and/or Aβ (**Fig. 5c & d**). In Alzheimer’s vaccine development, such antibodies must recognise pathological deposits within the brain, marking them for subsequent immune-mediated clearance. We therefore examined whether day-28 immune sera recognise pathology in post-mortem brain sections from transgenic models of Alzheimer’s disease expressing human pathogenic tau (TAU58/2) or Aβ (APP/PS1) (**Fig. 6**).

**Figure 6.**
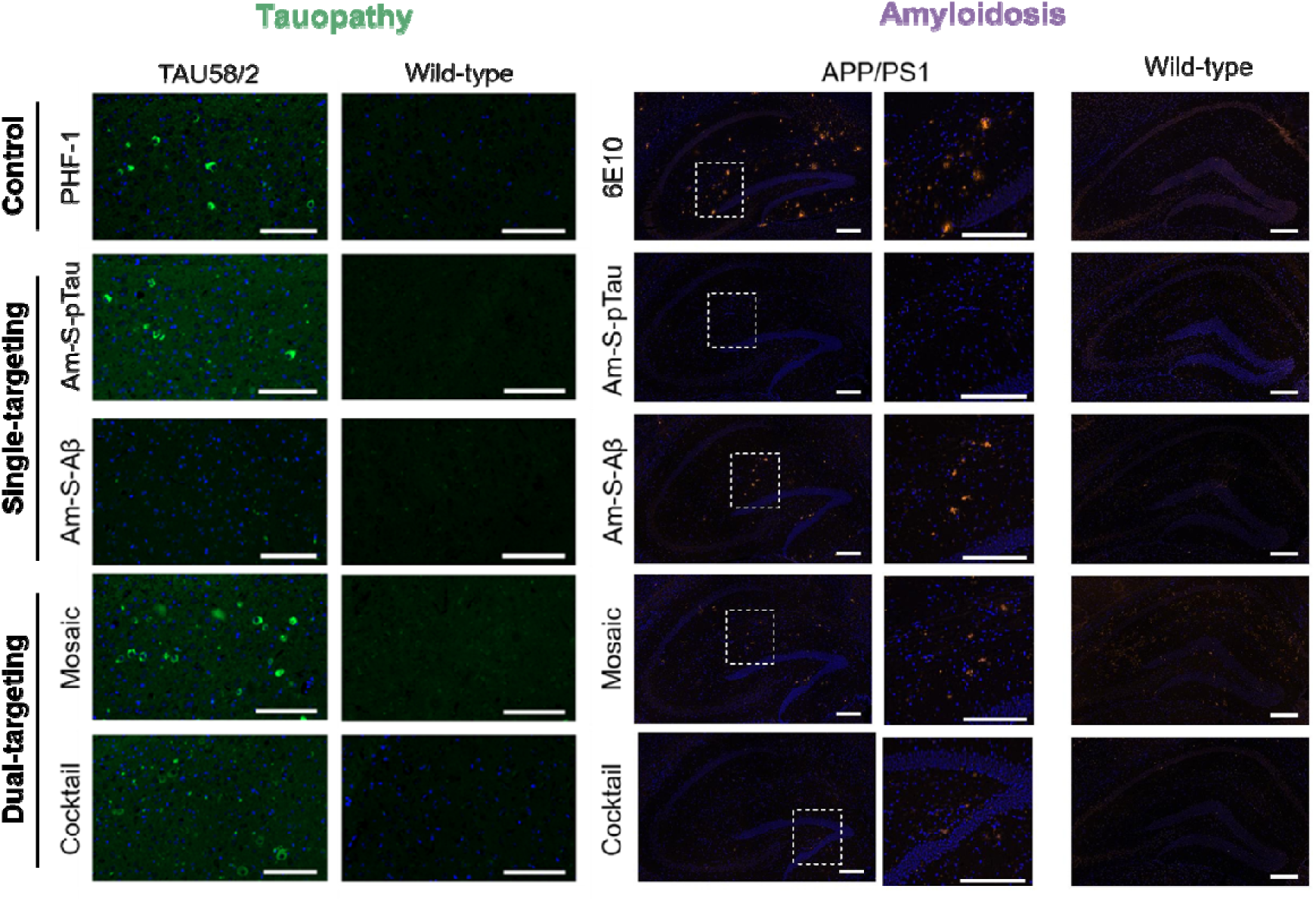
Immunoreactivity of nanovaccine-elicited antibodies on ex vivo brain tissue from transgenic AD mice. Representative immunohistochemical staining of **(Left panel)** amygdala sections from tauopathy TAU58/2 mice with the corresponding region from wild-type mice controls (*n* = 1); and **(Right panel)** hippocampal sections from amyloidogenic APP/PS1 mice with the corresponding region from wild-type controls (*n* = 1). As indicated, ex vivo brain sections were incubated with sera from C57BL/6J mice immunised with single-targeting nanovaccines (Am-S-pTau or Am-S-Aβ), or dual-targeting mosaic or cocktail formulations. Positive control antibodies were included: PHF-1 that recognizes pTau (pSer396/404); or 6E10 that binds Aβ (residues 1-16/17). Pathology-bound IgG was detected using Alexa Fluor 488 (green) and Alexa Fluor 568 or 647 (orange). Cell nuclei were counterstained with DAPI (blue). Scale bars = 200 µm (TAU58/2); or 500 µm (APP/PS1) and 200 µm (zoomed-in region, APP-PS1).

TAU58/2 mice express P301S mutant human tau and develop progressive tau hyperphosphorylation and neurofibrillary tangle pathology ^67^, with the amygdala representing a primary site of involvement ^68,69^. Immunohistochemical staining of ex vivo TAU58/2 amygdala sections incubated with sera from mice immunised with Am-S-pTau or mosaic nanocages enabled visualisation of intracellular antibody binding within neuronal structures in the posterior amygdala region (**Fig. 6, Left panel**). The spatial distribution was comparable to the pathogenic tau marked by the PHF-1 reference antibody. Sera from the cocktail group also showed detectable intra-neuronal binding, although staining appeared less frequent. No signal was apparent in amygdala sections incubated with sera from the Am-S-Aβ group or in the corresponding tissue from wild-type mice.

APP/PS1 mice develop human AD-like amyloid deposition in the hippocampus. Immunohistochemical staining of APP/PS1 hippocampal sections incubated with sera from mice immunised with Am-S-Aβ or mosaic nanocages revealed discrete extracellular Aβ plaque-like labelling (**Fig. 6, Right panel**). The sera-stained sections display compact centres and fibril-like extensions as expected of AB plaques, mirroring plaque formation observed with the 6E10 control antibody. Sera from the cocktail group also produced detectable plaque binding, but with lower apparent intensity, consistent with the reduced anti-Aβ titres measured in this group (**Fig. 5d**). Minimal signal was observed in sections exposed to sera from the Am-S-pTau group or in the wild-type tissue controls.

These data demonstrate that antibodies generated against pTau and/or Aβ antigens (co-)displayed on the Am-S nanoscaffold recognize and selectively bind the corresponding human pathological forms of tau and/or Aβ in situ. Future work should test the binding ability of the antibodies to human AD post-mortem tissue.

## Conclusions

We have identified a novel *T* = 1 encapsulin from *Alkaliphilus metalliredigens* (AmEnc) and characterized it as a classical Family 1 system that encapsulates FLPs, likely contributing to iron-mediated oxidative stress regulation in its host. Leveraging this encapsulin, we developed a SpyCatcher-decorated modular nanoscaffold called Aml7JS, capable of covalent conjugation to SpyTagged antigens. Am-S exhibited high-level soluble expression and favorable yields in *E. coli* (∼200 mg/L), representing a 4- to 10-fold increase over reported soluble yields for Spytag-decorated MxEnc (*T* = 3) ^21,44^ and TmEnc (*T* = 1) nanocages. Am-S nanocages retained structural integrity and solubility following multiple freeze-thaw cycles, storage from −80C°C to ambient temperatures, and lyophilization. Collectively, these properties establish Am-S as a robust and manufacturable nanoscaffold.

Future Am-S manufacturing should focus on process scalability, including optimization of upstream production via fed-batch or continuous fermentation and downstream purification using multi-step chromatography and tangential flow filtration ^6^. While bacterial expression systems offer high titers and are used to produce approximately 30% of VLPs ^70^, residual endotoxin remains a critical barrier to clinical translation of PNPs. Rigorous endotoxin removal strategies, or the use of engineered *E. coli* strains with disrupted lipopolysaccharide (LPS) biosynthesis, such as ClearColi™, an approach successfully applied in PNP production ^71^, provide practical routes to minimizing endotoxin burden and meeting regulatory requirements.

As a proof-of-concept for use in nanovaccine development, Am-S was employed to display AD antigens derived from Aβ and pTau. The modular Am-S platform enabled high-density, stoichiometrically controlled conjugation to generate both monovalent and dual-targeting mosaic nanocages. Following administration into C57BL/6J mice, Am-S was well-tolerated with no observed adverse effects, establishing an initial safety profile. Although encapsulin-based PNPs have not entered human clinical trials, they have demonstrated favourable biocompatibility across in vitro and in vivo models, including non-human primates ^15,17,20^. In vivo studies also demonstrated that antigen display on Am-S markedly enhanced immunogenicity relative to free antigen peptides, eliciting strong, Th2-biased IgG responses, a profile associated with reduced risk of neuroinflammation. Importantly, these dual-targeting nanocages generated anti-pTau and anti-Aβ antibodies capable of recognizing pathological human forms of both proteins in ex vivo brain tissue isolated from AD mice models, validating the platform’s capacity for multivalent targeting of self-antigens. Future studies should examine the Am-S platform’s long-term toxicity profile and immunogenicity.

Building on these results, the validated Aβ and pTau antigen combinations on Am-S could be used to immunise AD mice models (e.g., humanized AppNL-F/MAPT double knock-in mice ^72^), to evaluate the vaccine’s capacity to reduce pathological Aβ and pTau accumulation and to prevent or slow cognitive decline. Moreover, the modularity of the Am-S platform permits the attachment of other AD-specific target antigens, including alternative Aβ species (e.g., pyroglutamate-modified Aβ) and distinct tau phosphorylation sites (e.g., pSer202/pThr205, pThr231) ^73–75^. This flexibility enables systematic assessment of antigen combinations, conjugation density, and dosing regimens to account for the heterogeneity and temporal changes in Aβ/pTau pathology observed in human patients. Collectively, this study provides a practical framework for the development of multi-target active immunotherapies, which offer advantages over existing, passive antibody approaches and can address the complex pathology of AD by generating polyclonal, long-lasting responses against multiple synergistic targets.

Taken together, this work positions Am-S as a versatile scaffold for multivalent antigen display and nanovaccine development, supporting multi-target prophylactic and therapeutic vaccination strategies against complex diseases.

## Materials and methods

### Plasmid construction, protein expression and purification

#### A. metalliredigens-derived encapsulin nanocage (AmEnc)

The *E. coli* codon-optimised encapsulin gene from *A. metalliredigens* (WP_012065385.1) (**Table S2**) was synthesised as a gBlock gene fragment (Integrated DNA Technologies) with flanking NcoI and BamHI restriction sites and cloned into the pACYC-Duet-1 expression vector (Novagen, Merck, USA). The construct was transformed into *E. coli* BL21(DE3) competent cells (New England Biolabs, USA).

Freshly transformed cells harbouring the expression plasmid were grown aerobically at 37 °C in Luria Bertani (LB) broth (Sigma-Aldrich) supplemented with carbenicillin (100 µg/mL, Thermo Fisher Scientific). Protein expression was induced at OD600 0.5–0.6 by addition of 0.4 mM isopropyl-β-D-thiogalactopyranoside (IPTG) (Thermo Fisher Scientific), followed by incubation for 4 h at 37 °C. Cells were harvested by centrifugation (6,000 *g*, 10 min, 4 °C) and pellets were stored at −30 °C.

Pellets from a 1 L culture were resuspended in lysis buffer (50 mM HEPES, pH 7.4) and disrupted by three passages through a French pressure cell. Insoluble debris was removed by centrifugation (14,000 *g*, 10 min, 4 °C). The clarified soluble fraction was incubated at 75 °C for 20 min to denature host proteins and partially purify recombinant AmEnc nanocages, followed by centrifugation (14,000 *g*, 15 min, 4 °C). The supernatant was supplemented with PEG 8000 (100 mg/mL) and NaCl (20 mg/mL) and incubated on ice for 45 min. Precipitated proteins were collected by centrifugation (10,000 *g*, 10 min, 4 °C) and resuspended in 13 mL HEPES buffer. Samples were clarified using a 0.2 µm filter (Millipore). Next, size-exclusion chromatography (SEC) was performed using a HiPrep 26/60 Sephacryl S-500 HR column (Cytiva) equilibrated in 50 mM HEPES pH 7.4 buffer at 1.3 mL/min. Further purification was achieved by anion-exchange chromatography using a HiPrep 10/16 Q column (Cytiva) with a step elution gradient of 0–1 M NaCl in HEPES buffer. Purified AmEnc nanocages eluted at 300–400 mM NaCl. Fractions were analysed by PAGE, and selected pooled fractions were concentrated using Amicon Ultra-15 centrifugal filter units with a 100 KDa cut-off (Merck). The final preparation was adjusted to 1 mg/mL in 50 mM HEPES (pH 7.4), filtered through a 0.2 µm filter (Millipore), and stored at −30 °C for use in subsequent experiments.

#### SpyCatcher-decorated AmEnc (Am-S) nanoscaffold

The *E. coli* codon optimised AmEnc-SpyCatcher (Am-S) sequence (**Table S2**) was synthesised by Twist Bioscience into the pET-24(+) expression vector and transformed into *E. coli* BL21(DE3) competent cells (New England Biolabs). Cells from a single colony were cultured in a 5 mL starter LB broth culture containing 50 µg/mL of kanamycin (Thermo Fisher Scientific) and grown for 16–18 h at 37 °C with shaking at 200 rpm. This starter culture was then diluted into 500 mL of LB and kanamycin (50 µg/mL) and grown at 37 °C/200 rpm until reaching an optical density at 600 nm (OD_600_) of 0.5–0.6. Protein expression was induced with the addition of IPTG to a final concentration of 0.1 mM and incubated for 16–18 h at 20 °C/200 rpm. Cells were harvested via centrifugation (8,000 *g*, 4 °C, 15 min) and stored at −30 °C until further use.

Cells were resuspended (5 mL/g wet cell mass) in binding buffer (20 mM sodium phosphate pH 7.4, 500 mM sodium chloride (NaCl), 40 mM imidazole) supplemented with 25 U/mL benzonase nuclease, cOmplete mini EDTA-free protease inhibitor (1 tablet/30 mL), and 1 mM phenylmethanesulfonyl fluoride (PMSF) (Sigma-Aldrich), incubated on ice for 15 min, and lysed via direct sonication at 50% amplitude for 5 min on ice with 10/20 s on/off pulsation, repeated 4 times. Insoluble debris was removed via centrifugation (10,000 *g*, 4 °C, 15 min), and the soluble protein was filtrated using a 0.22 µM syringe filter (Sartorius). The clarified protein was loaded with a flow rate of 1 mL/min onto 2x connected 5 mL HisTrap HP columns (Cytiva) pre-equilibrated with binding buffer for IMAC. The bound protein was then washed with 50 column volumes (CV) of binding buffer supplemented with 0.1% v/v Triton X-114 (Thermo Fisher Scientific) for on-column endotoxin removal ^42^, followed by a wash of binding buffer (20 CV) to remove the Triton X-114 (5 mL/min). Bound Am-S was further washed at 100 mM imidazole (5 CV) to remove contaminating proteins before finally being eluted at 240 mM imidazole (5 CV). Fractions containing Am-S nanocages were then subject to SEC using a HiPrep 26/60 Sephacryl S-500 column (Cytiva) equilibrated with 20 mM sodium phosphate buffer pH 7.4 with 150 mM NaCl and eluted over 1.5 CV at 1 mL/min. Fractions containing Am-S were pooled and concentrated using 100 KDa Amicon Ultra-15 centrifugal filter units with a cut-off. All purification steps were conducted at 4 °C. Recombinant bacterial host DNA was confirmed to be <0.6 (A260/280) via nanodrop (Nanodrop One/OneC Microvolume UV-Vis Spectrophotometer, Thermo Fisher Scientific) and protein concentration was determined via Bradford assay as per manufacturer’s instructions (Pierce™ Bradford Protein Assay Kit, Thermo Fisher Scientific). Purified Am-S was temporarily stored at 4°C before further use.

Two purifications of 100 mL of culture from two independent 500 mL expressions done on different days were used to estimate yield, based on Bradford assays and SDS-PAGE densitometry.

### Conjugation reactions

The SpyTag peptide sequence was joined by a flexible linker to the pTau peptide double phosphorylated at sites Ser396 and Ser404, pTau_389–408_ (S-pTau), and to the Aβ_1-6_ peptide (S-Aβ) (**Table S2**). The linear peptides were synthesised using automated solid phase peptide synthesiser (Liberty Blue, CEM) based on Fmoc-chemistry using diisopropylcarbodiimide (DIC) and Oxyma as coupling reagent. Peptides were then subjected to global deprotection with TFA/TIPS/H2O (95/2.5/2.5) for 3 h and precipitated to cold diethyl ether. The resulting residue was lyophilised. Crude peptides were purified by semi-preparative HPLC using water and acetonitrile with 0.1% formic acid as mobile phase. Peptide purity was >95%, validated by HPLC and LC-MS.

The antigens were reconstituted in 1x phosphate-buffered saline (PBS). Am-S was conjugated with S-pTau at a 1:4 molar ratio, S-Aβ at a 1:3 molar ratio, and the mosaic formulation was conjugated with a 1:4:1 molar ratio of Am-S:S-pTau:S-Aβ. Conjugations were done at 4 °C for 16 h with gentle rotation. Amicon Ultra-15 centrifugal filter units (100 KDa) were used to buffer exchange conjugates into PBS and remove any excess unbound antigens prior to centrifugation (16,900 *g*, 30 min, 4 °C) to remove any potential insoluble aggregations. Am-S was freshly purified and conjugated with antigens the day prior to each injection.

### Polyacrylamide gel electrophoresis

#### Sodium dodecyl-sulfate polyacrylamide gel electrophoresis (SDS-PAGE)

A Bio-Rad mini-protean system (Bio-Rad laboratories) was used for SDS-PAGE analysis. Protein samples were mixed in a 1:1 ratio with 2x Laemmli Sample Buffer (BioRad) with 50 mM 1,4-dithiothreitol and heated at 99 °C for 10 min with shaking at 300 rpm in a ThermoMixer (Eppendorf). Electrophoresis was performed at 200 V for 35 min on 4−20% or 10% polyacrylamide gels (Mini-PROTEAN TGX, BioRad) in SDS running buffer (25 mM Tris, 192 mM glycine, 0.1% SDS, pH 8.3). Gels were stained following the Coomassie G-250 safe stain protocol ^76^ and molecular weights compared using a commercial protein marker (Precision Plus Protein Standards, BioRad). Densitometry was done using either ImageJ or Image Lab 6.1 (BioRad).

#### Blue native PAGE (BN-PAGE)

An XCell SureLock Mini-Cell Electrophoresis System (Thermo Fisher Scientific) was used for BN-PAGE analysis. Protein samples were mixed in a 1:4 ratio with 4x Native-PAGE sample buffer (Thermo Fisher Scientific), loaded into NativePAGE 3−12% Bis-Tris protein gels (Thermo Fisher Scientific), and ran with NativePAGE Cathode Buffer Additive and Running Buffer (Thermo Fisher Scientific). Electrophoresis was performed at 4 °C at 150 V for 1.5 h, followed by 250 V for 1 h. Gels were fixed in 40% methanol and 10% acetic acid and de-stained in 8% acetic acid. Molecular weights were compared using a commercial protein marker (NativeMark Unstained Protein Standard, Thermo Fisher Scientific).

### Dynamic light scattering (DLS)

DLS was performed on samples with a final concentration 0.15 mg/mL in PBS using a Malvern Zetasizer Nano ZS. Three measurements were performed at 25 °C using a plastic microcuvette (ZEN0040, Malvern). Data analysis was performed in Zetasizer Nano software.

### Transmission electron microscopy (TEM)

Samples (10 µL, 0.1 mg/mL) were deposited onto carbon film coated 300 mesh copper grids (ProSciTech) and negatively stained with uranyl acetate replacement stain (UAR-EMS) (1:4 dilution) for 1 h, washed with ultrapure water, and allowed to dry for at least 24 h. TEM was performed using a FEI Tecnai G2 20 operating at 200 kV.

### Cryo-EM sample preparation and data collection

#### AmEnc

Purified encapsulin protein (∼3 mg/mL) was applied (4 μL) to glow-discharged Quantifoil R2/2 copper grids (Quantifoil Micro Tools). Grids were blotted for 3 s at 4 °C and 90% relative humidity and plunge-frozen in liquid ethane using a Leica EM GP (Leica Microsystems).

Cryo-EM data were collected on a FEI Talos Arctica transmission electron microscope operated at 200 kV using EPU automated acquisition software. A total of 1,310 movies were recorded on a Falcon III direct electron detector in counting mode at 120,000× magnification, corresponding to a calibrated pixel size of 1.25 Å. Each movie consisted of 40 frames with a total dose of 40 e^⁻^/Å².

#### Am-S

Purified Am-S protein (2 mg/mL) was applied (4.5 μL) to glow-discharged Quantifoil R2/2 copper grids. Grids were blotted for 4.5 s at 4 °C and 90% relative humidity and plunge-frozen using a Leica EM GP.

Data were collected on a FEI Talos Arctica microscope operated at 200 kV with EPU software. A total of 2,310 movies were recorded on a Falcon III detector in counting mode at 150,000× magnification (pixel size 0.986 Å). Each movie consisted of 40 frames with a total dose of 30 e^⁻^ /Å².

### Image processing and 3D reconstruction

#### AmEnc

Movie frames were aligned using MotionCorr2 ^77^, and CTF parameters were estimated with Gctf ^78^. A total of 1,310 micrographs were selected for the processing in RELION 3.0 ^79^.

Particles (235,978) were automatically picked and extracted with a box size of 300 pixels. Following 2D classification, 100,449 particles were selected for 3D analysis. An initial model (EMD-3608) was used for 3D classification, yielding 77,899 particles for refinement.

3D refinement resulted in a map at 3.9 Å resolution, which was improved to 3.1 Å after CTF refinement, Bayesian polishing, and post-processing. Resolution was estimated using the gold-standard Fourier shell correlation (FSC) 0.143 criterion.

#### Am-S

All processing was performed in CryoSPARC v4.3.1 ^80^. Movie frames were motion-corrected, and CTF parameters were estimated using Patch CTF.

Particles were initially manually picked and subjected to 2D classification to generate templates for automated picking. After iterative 2D classification, 210,377 particles were selected for reconstruction.

An ab initio model with I2 symmetry was generated and used for non-uniform refinement. The final reconstruction reached a resolution of 2.58 Å based on the gold-standard FSC 0.143 criterion.

### Model building, refinement, validation, and data deposition

The initial models were derived from the previously published structure of *T. maritima* (PDB 7MU1) and refined against the experimental maps using PHENIX real-space refinement (v1.21.2-5419) ^81^. Model quality was assessed using PHENIX validation tools.

The cryo-EM density maps were deposited in the Electron Microscopy Data Bank (EMDB) under accession codes **EMD-68491 (AmEnc)** and **EMD-80547 (Am-S)**, and the corresponding atomic coordinates for AmEnc were deposited in the Protein Data Bank (PDB) under accession code **22NB.**

Molecular graphics and figures were prepared using UCFS Chimera v1.17 ^82^.

### Storage stability

#### Freeze-thaw stability of Am-S

To test the freeze-thaw storage stability of Am-S, 20 µL aliquots in Eppendorf tubes (2 mg/mL in PBS) were subject to 1 or 4 freeze-thaw cycles by storing at −80 °C until completely frozen, followed by thawing in water at ambient temperature. Samples were centrifuged (16,900 *g*, 30 min, 4 °C) to remove potential insoluble aggregations prior to DLS and SDS-PAGE densitometry analysis. Samples that had not been frozen were defined as 100% soluble.

#### Longer-term storage stability of Am-S

Longer-term storage in a range of temperatures was tested using 20 µL aliquots of Am-S in Eppendorf tubes (2 mg/mL in PBS) were stored at ambient temperature, 4 °C, −20 °C, and −80 °C for 6 weeks. Samples were analysed as above, and samples stored at −80 °C were defined as 100% soluble.

#### Lyophilisation storage stability of Am-S

To test Am-S stability after lyophilisation, 60 µL aliquots in Eppendorf tubes (2.5 mg/mL in PBS) were frozen at −80 °C and then lyophilised for 4 h using an Alpha 2-4 LDplus freeze dryer (Martin Christ) at 0.1 mbar and −80 °C. Samples were stored at ambient temperature overnight before reconstitution to the original volume with ultra-pure water and analysed as above, with sample collected prior to lyophilisation defined as 100% soluble.

### C57BL/6J immunisation

Male C57BL/6J mice were housed in a 12 h light-dark cycle-controlled facility with food and water available ad libitum. Animal experiments were approved by the University of Technology Sydney’s Animal Care and Ethics Committee (ETH24-9430) and performed according to the Australian Code for the Care and Use of Animals for Scientific Purposes, 8th Edition, 2013 guidelines. Mice were injected subcutaneously with 100 µL of formulation, with the concentration of vaccines adjusted to keep the amount of pTau/Aβ antigens consistent between groups, i.e., groups received either the Am-S-pTau vaccine (100 µg Am-S/dose), Am-S-Aβ vaccine (100 µg Am-S/dose), mosaic vaccine (200 µg Am-S/dose), or a cocktail mixture of both Am-S-pTau and Am-S-Aβ vaccines (100 µg + 100 µg Am-S/dose). Assuming complete conjugation, each dose contained 9 µg of pTau and/or 5.5 µg of Aβ. Adjuvants were mixed at a volume ratio of 1:1 as per manufacturer’s instructions immediately before administration. Mice were euthanised via cardiac puncture under isoflurane anaesthesia. Whole blood was allowed to clot for 30–60 min at ambient temperature before centrifugation (1,500 *g* for 10 min at 4 °C), and serum was collected and stored at −20 °C before analysis.

#### Preliminary assessment of Am-S-pTau immunogenicity

Mice (*n* = 4, 6–8 weeks old) were injected with the Am-S-pTau vaccine or controls followed by booster injections on day 14. Adjuvanted groups received injections supplemented with complete Freund’s adjuvant (FA) for the prime injection, and incomplete FA for the booster injection (Sigma-Aldrich). Mice were euthanised on day 21.

#### Immunogenicity of Am-S-pTau, Am-S-Aβ, mosaic, and cocktail vaccines

Mice (*n* = 4, 6–8 weeks old) were injected with vaccines or controls supplemented with AddaVax™ (ADV) or Alhydrogel^®^ (ALH) (Jomar Life Research). Booster injections were administered on day 14 and day 28. Blood was collected weekly via the submandibular vein before mice were euthanised on day 35.

### Enzyme-linked immunosorbent assay (ELISA)

Anti-pTau and anti-Aβ total IgG in mouse sera was determined by coating 384-well Nunc MaxiSorp Immuno Plates (Thermo Fisher Scientific) with 0.125 µg/well of either the pTau_389-408_ (pS396/pS404) peptide or the human β-Amyloid (1–42) peptide (Genscript, RP10017) overnight at 4 °C. Wells were washed 3x with 100 µL of PBST buffer (PBS + 0.05% Tween 20), blocked with 10% foetal bovine serum for 1 h at ambient temperature, and washed again. Serial two-fold dilutions of sera (starting from 1:128 in PBS) were added to the wells and incubated for 1 h at ambient temperature followed by washing as before. HRP-conjugated goat anti-mouse IgG (Sigma-Aldrich) was used as the secondary antibody, diluted 1:5,000 in PBST and incubated for 1 h at ambient temperature, followed by washing. TMB Chromogen Solution (for ELISA) (Thermo Fisher Scientific) was used as per manufacturer’s instructions and the reaction stopped with 6% phosphoric acid. Absorbance was measured at 450 nm with subtraction at 570 nm for optical correction using a Tecan M200 plate reader (Tecan, Switzerland). Antibody titre was defined as the highest dilution at which the absorbance was at least 3x the absorbance of equally diluted control sera from mice injected with adjuvant only.

For isotyping, ELISA’s were done as above using HRP-conjugated goat anti-mouse IgG1, IgG2c, or IgG2b secondary antibodies (Abcam) diluted 1:75,000.

### Immunohistochemistry

TAU58/2 mice and their non-transgenic (non-tg) littermates at 8 months old were anaesthetised and transcardially perfused with cold PBS (ethics approval number ARA 2017/053; donated by the Dementia Research Centre). One brain hemisphere was immersion-fixed in 4% paraformaldehyde, processed in an Excelsior tissue processor (Thermo Fisher Scientific), embedded in paraffin, and sectioned coronally at 3 µm. Brain sections from 9-month-old APP/PS1 transgenic mice showing pathological Aβ plaque accumulation (ethics approval number ETH19-4326; donated by Associate Professor Kristine McGrath) were paraffin embedded and sagittally sectioned at 5 µm.

Sections underwent antigen retrieval in EDTA buffer for 15 mins in a pressure cooker, were blocked in protein block for 2 h (0.1 M phosphate buffer, normal goat serum, BSA, Triton X-100), then incubated either with pooled (*n* = 2-4) serum collected at day 28 from vaccinated mice (diluted 1:10 to stain Tau58/2 brains and 1:3 to stain APP/PS1 brains, in protein block) or positive control antibodies (PHF1 – 1:250; 6E10 – 1:250) overnight at 4 °C. After washing, the slides were incubated for 2 h at room temperature, protected from light, with DAPI (1:500, ThermoFisher Scientific) and fluorescent secondary antibody (goat anti-mouse IgG AF488 or AF568, Invitrogen; or goat anti-rabbit IgG AF647, Invitrogen for binding with 6E10) cocktail, washed as previous, cover slipped, then imaged using the Zeiss Axioscan high throughput slide scanner, or the Nikon TiE2.

### Statistical analysis

Statistical analyses were done using the GraphPad Prism version 10.2.2 for Windows (GraphPad Software, Boston, Massachusetts USA, https://www.graphpad.com).

## Acknowledgements

I.B. was supported by a Dementia Australia Research Foundation (DARF) PhD scholarship. This work was also supported by the National Health & Medical Research Council (NHMRC, 2037822), DARF, the Mason Foundation, and the National Foundation for Medical Research & Innovation (NFMRI). We thank Dr Daryl Ariawan (MQ Dementia Research Centre, NSW, Australia) for peptide synthesis; A/Prof Kristine McGrath (UTS, NSW, Australia) for provision of APP/PS1 transgenic mouse brain tissue; and the late Dr Peter Davies (Albert Einstein College of Medicine, NY, USA) for the PHF-1 antibody, provided through The Feinstein Institutes for Medical Research (NY, USA). J.R thanks Dr. Fumiaki Makino (Osaka University, Osaka, Japan) for his initial modelling assistance with AmEnc. We acknowledge the use of equipment and infrastructure within the Microbial Imaging Facility and the Ernst Facility, Faculty of Science (UTS), and the Electron Microscope Unit within the Mark Wainwright Analytical Centre (UNSW, NSW, Australia). Molecular graphics and analyses were performed using UCSF ChimeraX, developed by the Resource for Biocomputing, Visualization, and Informatics at the University of California, San Francisco, with support from National Institutes of Health grant R01-GM129325 and the Office of Cyber Infrastructure and Computational Biology, National Institute of Allergy and Infectious Diseases.

We acknowledge and pay respect to the Gadigal people, the traditional custodians of the land on which this research was conducted.

