## Supplementary Materials for "Engineering a Novel Bacterial Encapsulin for Programmable Surface Functionalization: From Single-Target to Mosaic Nanovaccines"

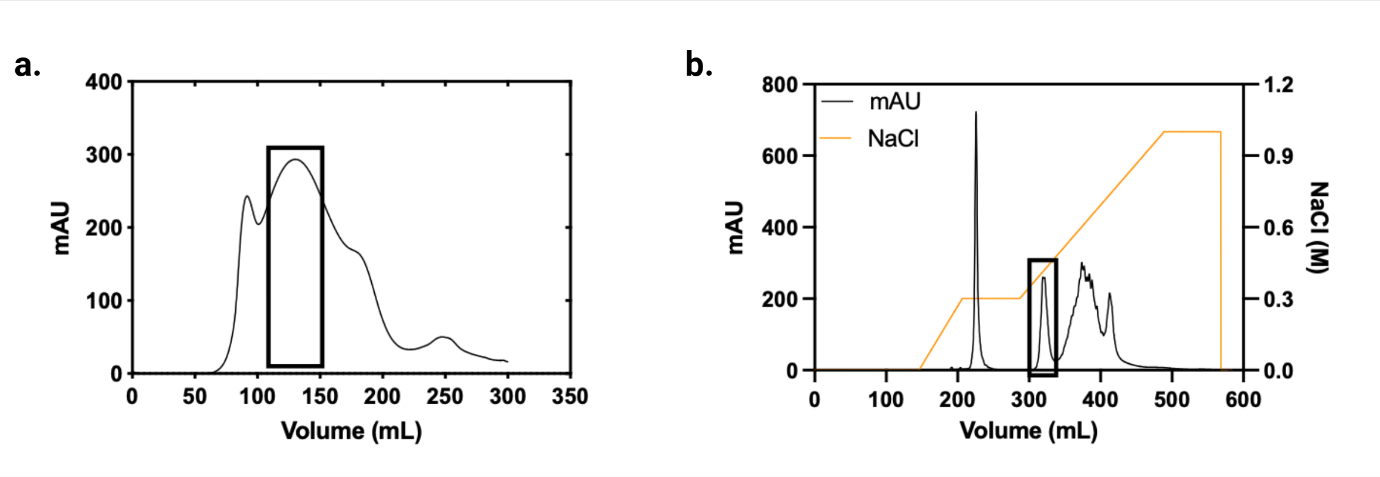

**Supplementary Figure S1. Example chromatograms of AmEnc purifications.** (**a**) Size Exclusion Chromatography (SEC). (**b**) Anion Exchange Chromatography of SEC purified AmEnc. The peaks corresponding to AmEnc are highlighted in black squares.

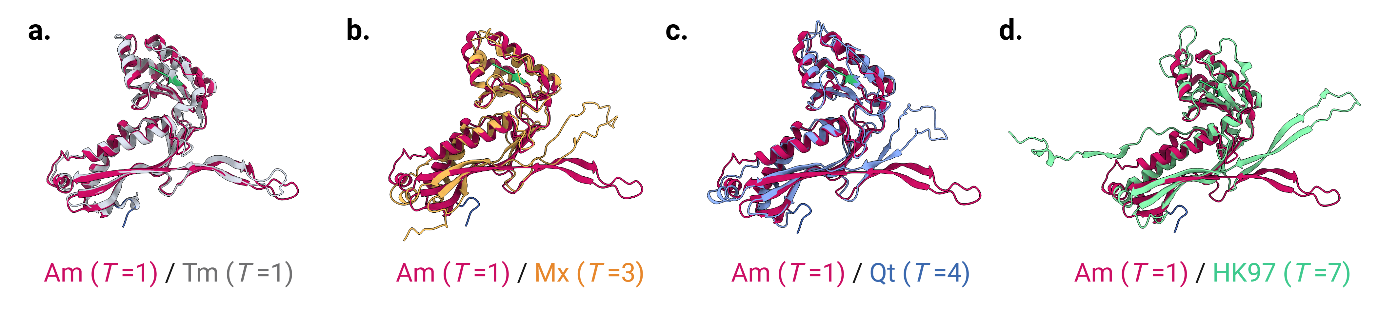

**Supplementary Figure S2. Structural superposition of the AmEnc subunit (pink) with previously characterized encapsulin subunits and the ancestral HK97 bacteriophage capsid protein.** Comparisons include: (**a**) *T*=1 TmEnc (PDB: 3DKT; grey), (**b**) *T*=3 MxEnc (PDB: 4PT2; orange), (**c**) *T*=4 QtEnc (PDB: 6NJ8; blue), and (**d**) *T*=7 HK97 bacteriophage (PDB: 1OHG; green). The AmEnc subunit is most structurally similar to the *T*=1 TmEnc scaffold.

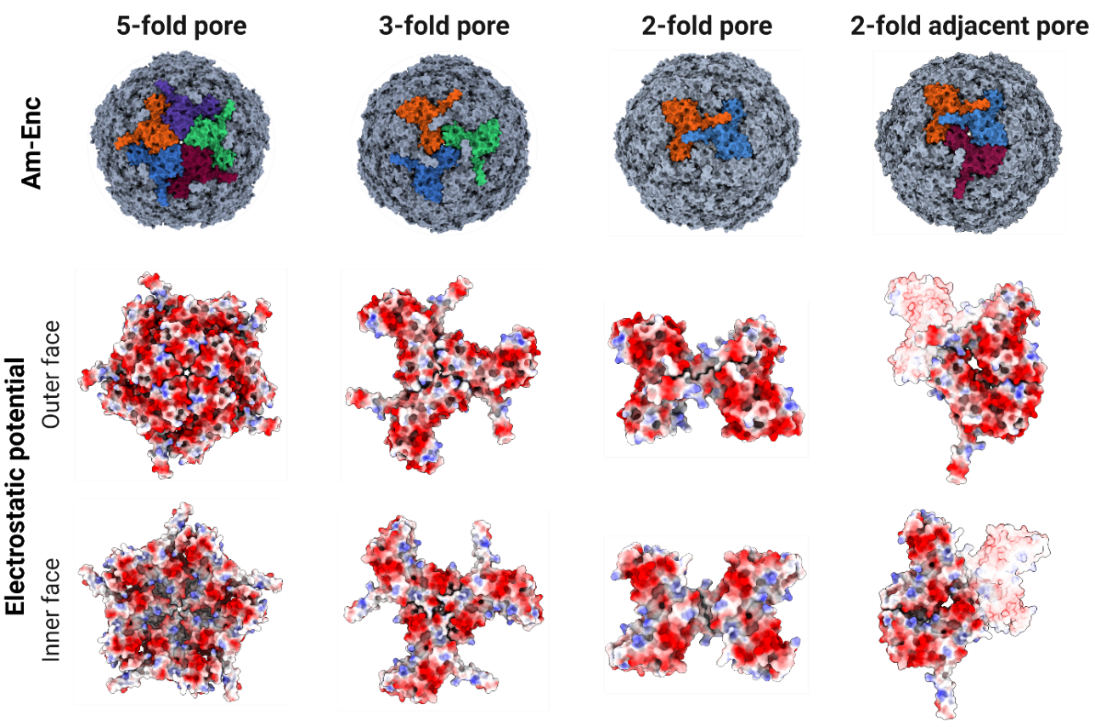
 Supplementary Figure S3. AmEnc nanocage surface pores and electrostatic potentials. (Upper panel) Surface density maps generated in ChimeraX displaying the four primary pores located at the 5-fold, 3-fold, 2-fold, and 2-fold adjacent symmetry axes. Individual subunits are uniquely colored to highlight the respective symmetry axes. (Lower panel) Exterior (outer) and luminal (inner) views of the symmetry pores colored by electrostatic potential. Negative and positive potentials are shown in red and blue, respectively.

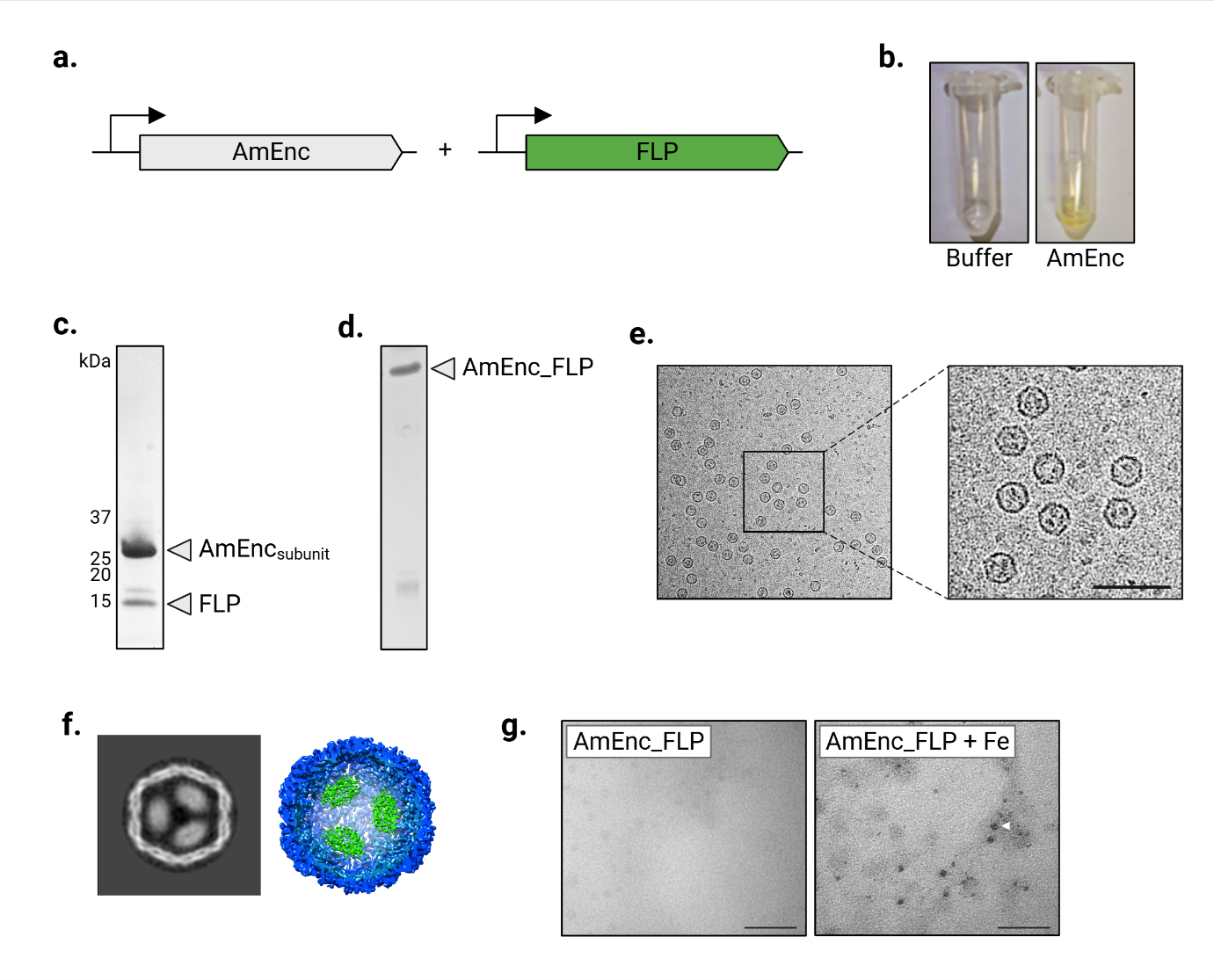

**Supplementary Figure S4. Structural and functional characterization of AmEnc and its native cargo.** (**a**) Schematic of the co-expression system encoding the AmEnc shell and the ferritin-like protein (FLP) cargo, which also comprises a C-terminal encapsulation signal (Esig) peptide that mediates selective cargo recruitment into the assembling encapsulin shell. (**b**) Visual confirmation of purified AmEnc displaying a vibrant yellow coloration compared to 50 mM HEPES buffer, indicative of surface-bound flavin cofactors. (**c**) SDS-PAGE of the co-purified complex, showing bands for the AmEnc subunit (29.9 kDa) and the ESig-tagged FLP cargo (13.3 kDa). (**d**) Native-PAGE verifying the self-assembly and integrity of the Enc_FLP complex. (**e**) Representative cryo-EM micrographs of assembled FLP-loaded AmEnc nanocage complexes (Enc_FLP); the encapsulated cargo is visible as discrete internal densities within the shells. Scale bar = 50 nm (**f**) Cryo-EM 2D class average (**left**) and 3D density map (**right**) of the Enc_FLP complex at 6 Å resolution. The shell is shown in blue and the encapsulated multimeric FLP cargo in green, revealing organized cargo density within the lumen. (**g**) TEM micrographs demonstrating the iron-mineralization functionality of Enc_FLP before (**left**) and after (**right**) exposure to aqueous ferrous iron; the latter displays distinct electron-dense mineralized iron cores. Scale bars = 100 nm.

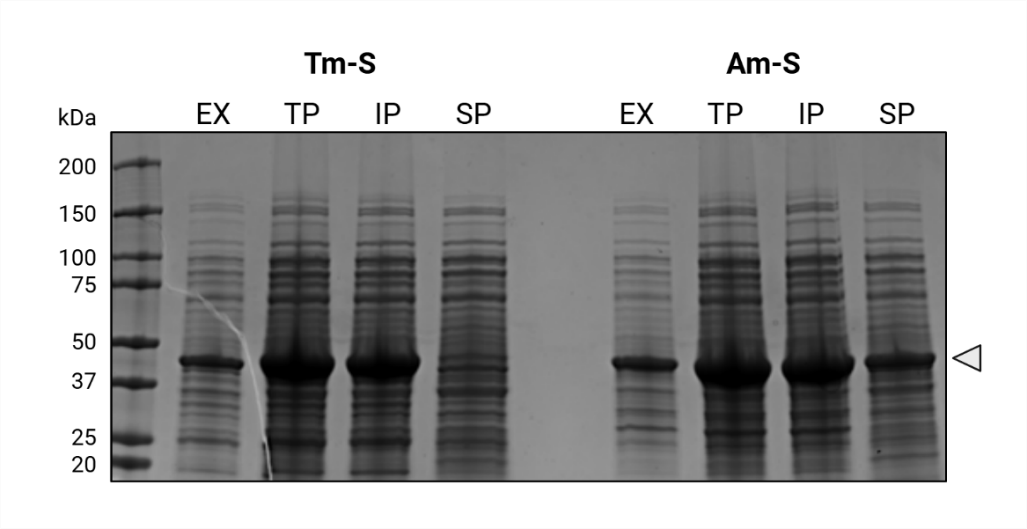

**Supplementary Figure S5.** Coomassie-stained SDS-PAGE gel used to visualise the expression (EX) and total protein (TP), insoluble protein (IP), and soluble protein (SP) fractions of Tm-S and Am-S after cell lysis. 500 mL cultures were expressed, with 100 mL harvested and lysed for this comparison. TP and IP samples were diluted 1:4 prior to loading onto gel.

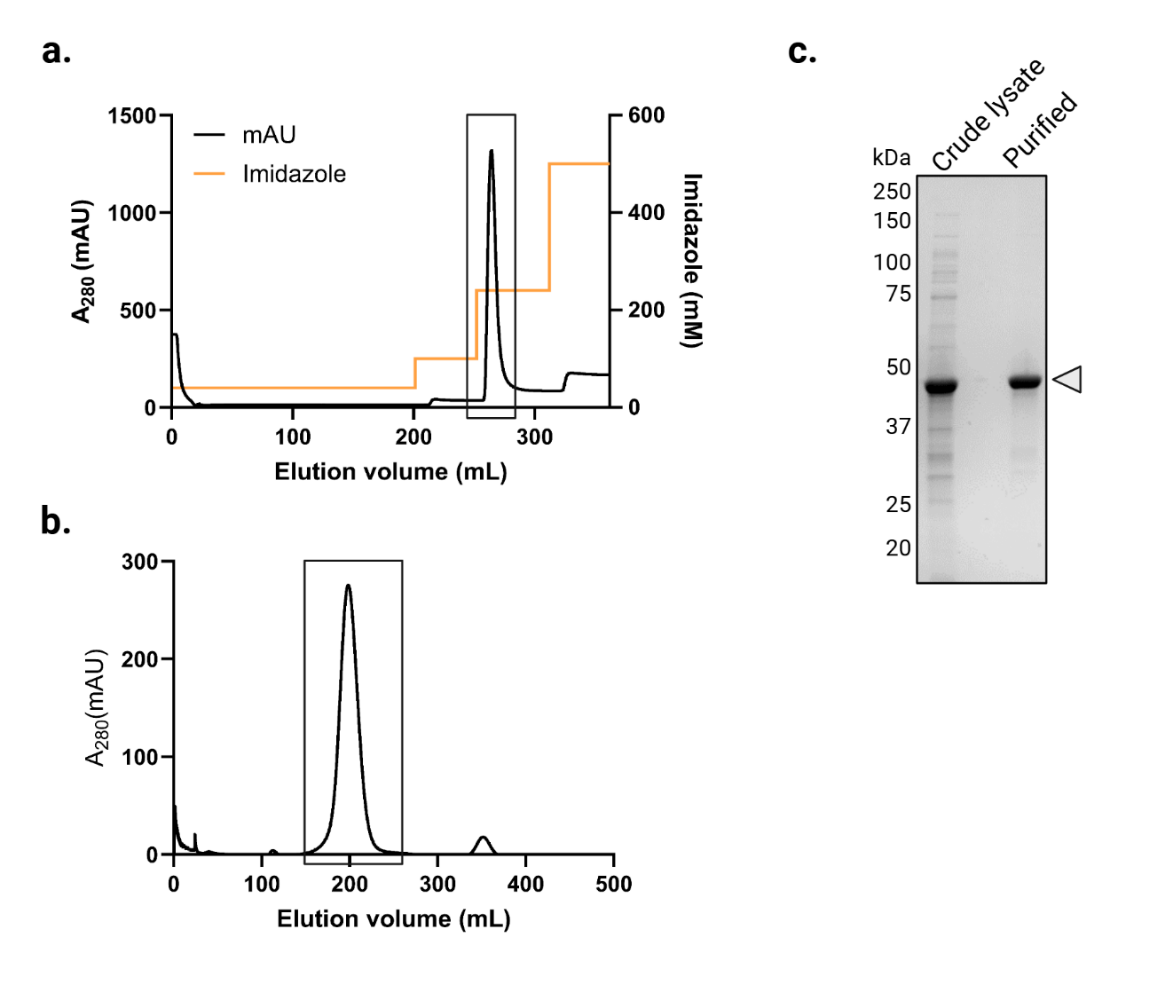

**Supplementary Figure S6. Purification of Am-S.** Representative chromatogram of (**a**) Immobilized Metal Affinity Chromatography (IMAC) and (**b**) SEC purification. The peaks corresponding to Am-S are highlighted in black squares. (**c**) Coomassie-stained SDS-PAGE showing Am-S before and after final purification.

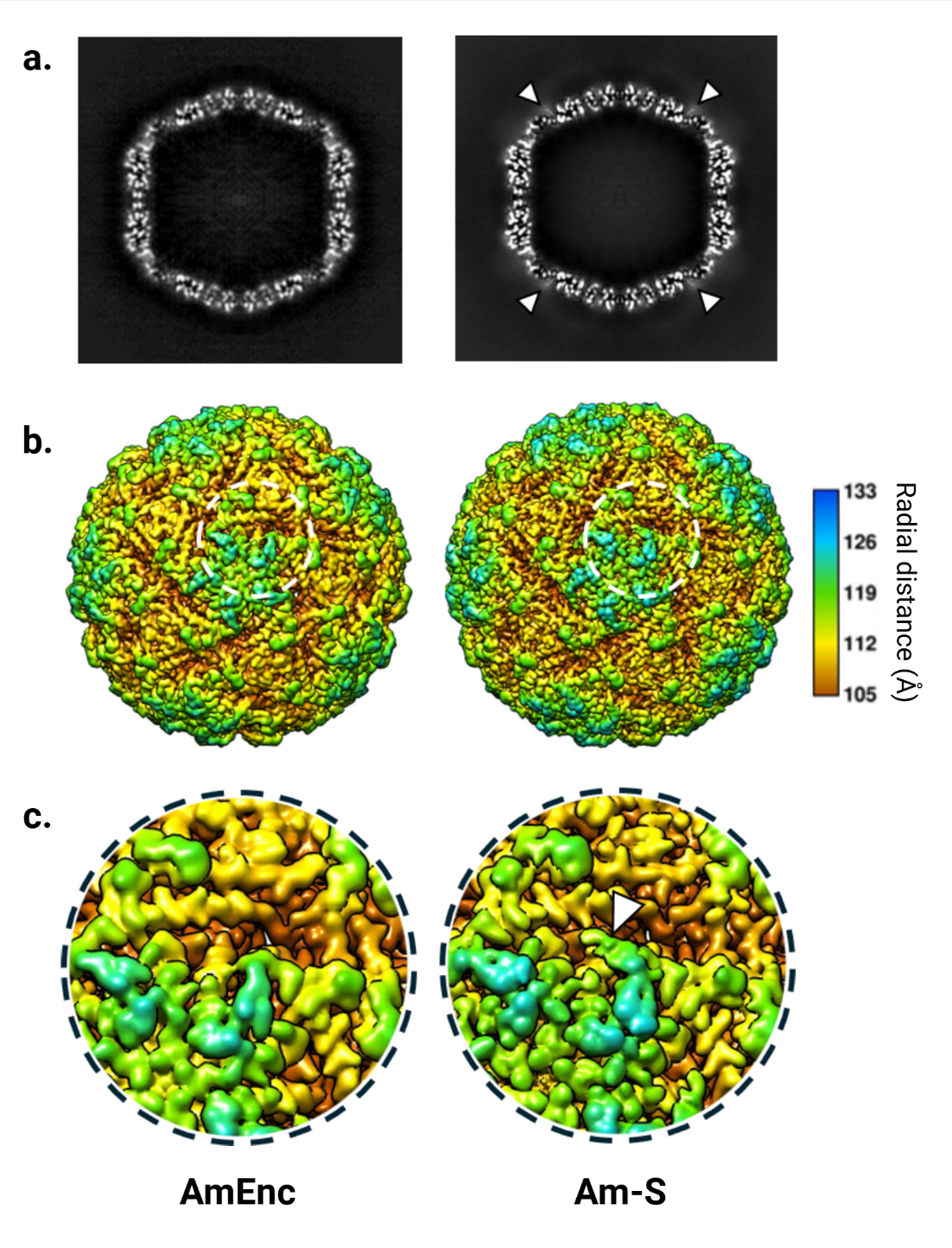

**Supplementary Figure S7.** (**a**) Central slices of the 3D density maps along the 2-fold axis for AmEnc and Am-S. (**b**) Corresponding 3D reconstructions rendered at the same threshold. (**c**) Enlarged views of the densities highlighted by white dashed circles in (b). The additional diffuse density, indicated by white arrowheads in (a), corresponds to the well-defined SC domain connected to the C-terminus. This additional density is also clearly visible in the 3D reconstructions in (b), where a representative position is marked by a white arrowhead in (c).

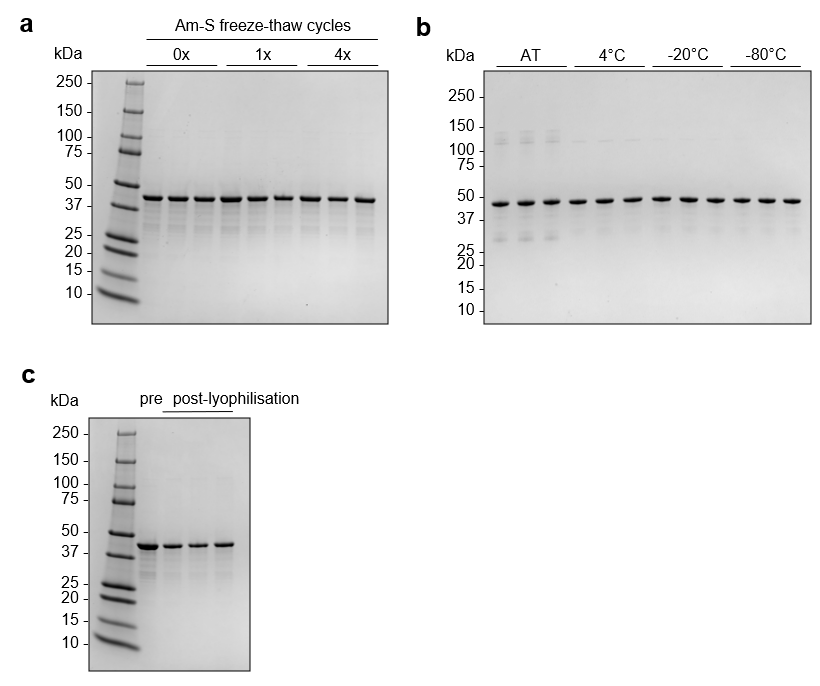

**Supplementary Figure S8. Storage stability of Am-S platform.** Coomassie-stained SDS-PAGE gels used to visualise any protein degradation and calculate solubility via densitometry for (**a**) Am-S after 1 and 4x freeze-thaw cycles, (**b**) Am-S after storage at the indicated temperatures for 6 weeks, and (**c**) Am-S before and after lyophilisation and storage at ambient temperature for 1 day. AT, ambient temperature.

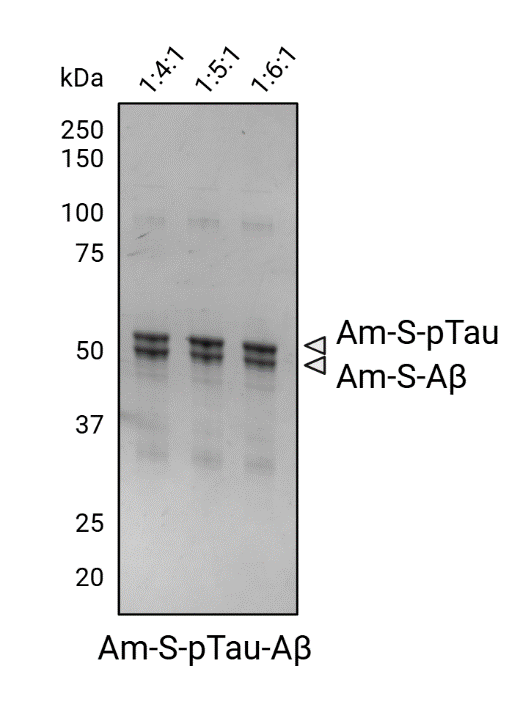

**Supplementary Figure S9.** Coomassie-stained SDS-PAGE gel used to visualise optimal molar ratios for conjugation of Am-S with S-pTau and S-Aβ. Gel densitometry indicated the 1:4:1 ratio resulted in the most even proportions of Am-S subunits conjugated to each antigen, with band % 49.2 and 50.8 for S-pTau and S-Aβ, respectively. The 1:5:1 ratio had a band % of 62.7/37.3 and the 1:6:1 ratio a band % of 63.4/36.6 for S-pTau/S-Aβ.

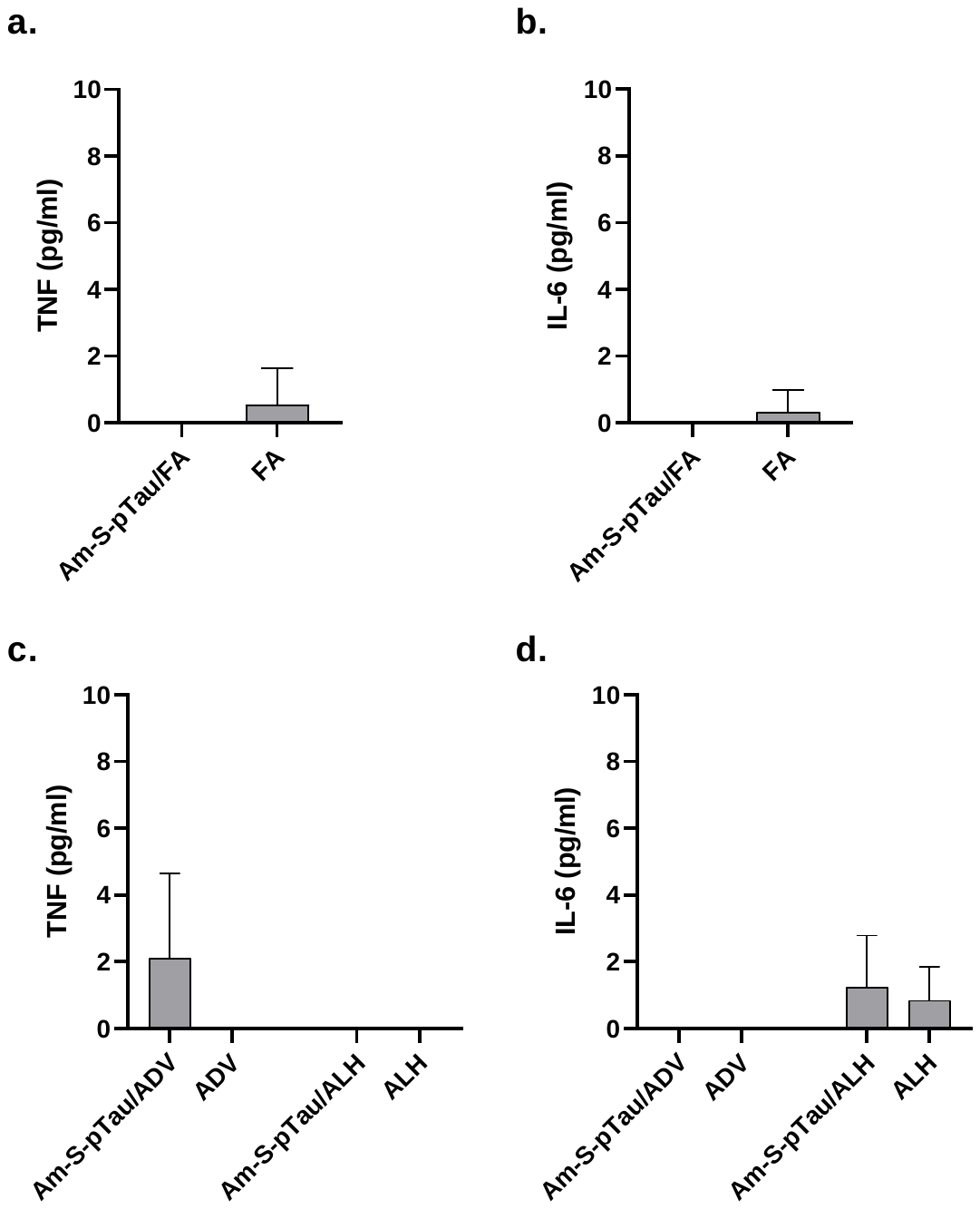

**Supplementary Figure S10.** Systemic markers of inflammation in sera collected at the experimental end point. Serum collected from mice vaccinated with Am-S-pTau/FA and FA only were assayed via cytometric bead array. Levels of (**a**) TNF and (**b**) IL-6 were below the limit of detection for the assay, and below physiologically relevant concentrations. Sera from mice vaccinated against Am-S-pTau formulated with both ALH and ADV adjuvants, and adjuvant only controls, similarly showed negligible levels of pro-inflammatory markers; (**c**) TNF, (**d**) IL-6. TNF – tumor necrosis factor; IL-6 – interleukin 6. Data are mean ± SD.

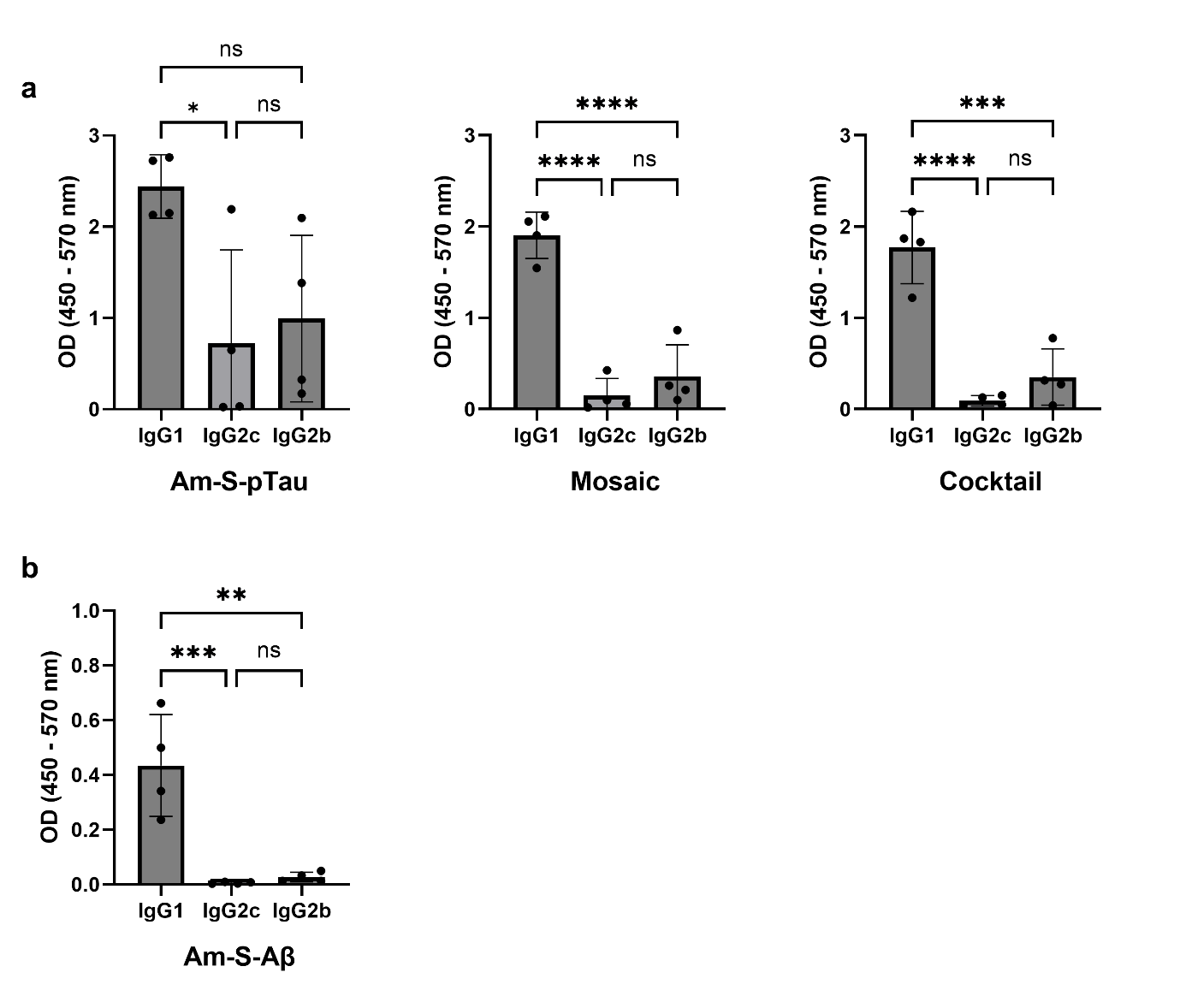

**Figure S11. Isotype profile of antibodies induced by vaccine variants.** (**a**) Anti-pTau antibody isotypes induced by vaccine variants. (**b**) Anti-Aβ antibody isotypes induced by Am-S-Aβ. Immune sera were diluted at 1:1000. Results are shown as mean ± SD with each replicate plotted. Data was analysed using a one-way ANOVA with Tukey’s multiple comparisons test. *****P* <0.0001; ****P* <0.001; ***P* = <0.01; **P* <0.05; ns (non-significant) >0.05.

**Supplementary Table S1.** Studies utilizing encapsulins as antigen carrier platforms.

| Enc | Antigen attachment | Application | Antigen | Adjuvant | Model | Admin route | Immune response | Ref |
| --- | --- | --- | --- | --- | --- | --- | --- | --- |
| TmEnc (*T =* 1) | Genetic fusion | Cancer vaccine | OT-1 peptide from ovalbumin protein. 8 AA | Poly(I:C) | C57BL/6 or OT-1 transgenic mice challenged with B16-OVA melanoma cells | SC (C57BL/6) or IP (transgenic) | Activated antigen-specific cytotoxic CD8^+^ T cells in both C57BL/6 and transgenic mice, and supressed tumour growth both therapeutically and prophylactically in challenged mice | ^1^ |
|  |  | Epstein-Barr virus vaccine (EBV) | EBV glycoprotein 350/220 (gp350) domain. 425 AA | Sigma adjuvant system | BALB/c mice and cynomolgus macaques | IM | Enhanced gp350-specific antibodies and neutralisation activity in both mice and macaques | ^2^ |
|  |  | Influenza vaccine | Matrix protein 2 ectodomain (M2e) epitope of influenza A virus H1N1 (9 AA). Also loaded green fluorescent protein (GFP) into interior | Freund’s adjuvant | BALB/c mice | SC | Generated antibody response against both M2e and GFP | ^3^ |
|  |  | Rotavirus vaccine | Core region of rotavirus VP8 protein (VP8*). 159 AA, 18 kDa | Alhydrogel (Alum hydroxide) | BALB/c mice | IM | Increased VP8*-specific antibodies and rotavirus neutralising antibodies | ^4^ |
|  |  | Poultry *Salmonella* vaccine | Ferric enterobactin receptor (FepA), extracellular loop 2 (E-L2), 28 AA | Freund’s adjuvant | BALB/c mice | SC | Elicited higher protein-specific antibodies as determined via ELISA (OD) compared with cage-only groups | ^5^ |
|  | SpyCatcher/SpyTag  (Enc fused with SpyTag) | HIV-1 vaccine primer | Fusion peptide (FP) site of vulnerability | Adjuplex | Rhesus macaques | Not reported | Priming with FP-conjugated Enc reduced non-neutralising trimer-base response and enhanced neutralisation | ^6^ |
|  | SpyCatcher/SpyTag  (Enc fused with SpyCatcher) | Lassa virus vaccine | Trimeric type 1-fusion glycoprotein complex (GPC). ~200 kDa | Adjuplex | Hartley guinea pigs and rhesus macaques | Not reported | Elicited neutralising response | ^7^ |
|  |  | SARS-CoV-2 vaccine | Receptor-binding domain (RBD) from WA1 and BA.5 strains. 473 AA | Sigma adjuvant system | BALB/c mice | IM | Generated high neutralisation titres | ^8^ |
|  |  | African Swine Fever virus (ASFV) vaccine | ASFV C129R protein, 129 AA | Freund’s adjuvant | BALB/c mice | IM | Increased anti-C129R humoral response and cellular response compared to antigen-only group | ^9^ |
|  |  |  | ASFV p54 protein, ~25 kDa | Freund’s adjuvant | BALB/c mice | SC | Enhanced p54-specific humoral response and cellular response compared to antigen-only group | ^10^ |
|  |  |  | ASFV p30 protein, 194 AA | Freund’s adjuvant | BALB/c mice | SC | Increased p30-specific humoral response and cellular response compared to antigen-only group | ^11^ |
|  | Intein-mediated transplicing | Antibody production pipeline | Antigens from ubiquitin-specific protease 24 (141 AA), murine cytomegalovirus protein M44 (97 AA), and immediate early protein 1 (131 AA) | N/A | BALB/c mice | IP | Elicited higher protein-specific antibody titres compared with antigen-only groups | ^12^ |
| MxEnc (*T* = 3) | Genetic fusion | SARS-CoV-2 vaccine | RBD peptide with K417N mutation. 20 AA | Engineered MxEnc to package bacterial single-stranded RNA from production host to enhance adjuvanticity | BALB/c mice | SC | Generated antibodies against both full-length and peptide RBD | ^13^ |
|  |  | Influenza vaccine | Influenza stem domain (pH1HA10). 140 AA | Sepivac SWE | BALB/c mice | IM | Elicited stem binding titres with protective immune response | ^14^ |
|  | SpyCatcher/SpyTag  (Enc fused with SpyTag) | SARS-CoV-2 vaccine | Monomeric RBD derivative (mRBD). 201 AA | Sepivac SWE | BALB/c mice | IM | Induced high mRBD and spike-binding titres with neutralising activity | ^15^ |

SC = Subcutaneous; IP = Intraperitoneally; IM = Intramuscular.

**Supplementary Table S2.** Sequences of constructs used in this study.

| Construct | Sequence |
| --- | --- |
| AmEnc | MDILKRDMAPLTESVWEEIDQRAAEVLKTHLSARRVVNIVGPKGWDYTVVPEGRLKKIEDNPGNVCTGMYQVKPLVEARISFKLDRWEMDNLIRGAKDIKLDALEEAAEKMAIFEENMLYNGYKPGDIEGLIEASSHKLSQFGNNGEEIMENLAQGMILLKEAYVDQPVTLVVGIDAWKRINREMQGHPLINRIQELTGSKVIYSPVVEGALLLPYDHEDLELTIGRDFSIGYEYHDAKTVQLFITESLTFRALNPDIIVVYNI |
| Am-S | MDILKRDMAPLTESVWEEIDQRAAEVLKTHLSARRVVNIVGPKGWDYTVVPEGRLKKIEDNPGNVCTGMYQVKPLVEARISFKLDRWEMDNLIRGAKDIKLDALEEAAEKMAIFEENMLYNGYKPGDIEGLIEASSHKLSQFGNNGEEIMENLAQGMILLKEAYVDQPVTLVVGIDAWKRINREMQGHPLINRIQELTGSKVIYSPVVEGALLLPYDHEDLELTIGRDFSIGYEYHDAKTVQLFITESLTFRALNPDIIVVYNIGGGSGGGSHHHHHHGGGSAMVDTLSGLSSEQGQSGDMTIEEDSATHIKFSKRDEDGKELAGATMELRDSSGKTISTWISDGQVKDFYLYPGKYTFVETAAPDGYEVATAITFTVNEQGQVTVNGKATKGDAHI |
| S-pTau_389-408_ pS396/pS404 (S-pTau) | AHIVMVDAYKPTKGSGSGAEIVYK(pS)PVVSGDT(pS)PRHL |
| S-Aβ_1-6_ (S-Aβ) | AHIVMVDAYKPTKGSGSDAEFRH |

*Phosphorylated amino acid: (pS)*

**Supplementary Table S3.** Cryo-EM data collection, refinement and validation statistics

|  | *AmEnc*  (EMD-68491)  (PDB 22NB) | *Am-S*  (EMD-80547) |
| --- | --- | --- |
| Data collection and Processing |  |  |
| Microscope | Talos Arctica | Talos Arctica |
| Detector | Falcon 3EC | Falcon 3EC |
| Magnification | 120,000 | 150,000 |
| Voltage (kV) | 200 | 200 |
| Electron exposure (e^-^/Å^2^) | 40 | 30 |
| Defocus range ($\boldsymbol{\mu}$m) | -1.5, -2.0, -2.5 | -1.5, -2.0 -2.5 |
| Pixel size (Å/pixel) | 1.25 | 0.986 |
| Micrographs collected (no.) | 1,310 | 2,310 |
| Initially particles (no.) | 235,978 | 216,700 |
| Final particles (no.) | 77,899 | 210,377 |
| Map resolution (Å) | 3.1 | 2.58 |
| FSC threshold | 0.143 | 0.143 |
| Axial symmetry | I2 | I2 |
| Refinement |  |  |
| Model resolution (Å) | 3.3 |  |
| FSC threshold | 0.5 |  |
| Correlation Coefficient (mask) | 0.85 |  |
| Map sharpening *B* factor (Å^2^) |  |  |
| Correlation Coefficient (volume) | 0.75 |  |
| Model composition |  |  |
| Non-hydrogen atoms | 2108 |  |
| Protein residues | 264 |  |
| Ligands | 0 |  |
| R.m.s. deviations |  |  |
| Bond lengths (Å) | 0.002 |  |
| Bond angels (^o^) | 0.572 |  |
| Validation |  |  |
| MolProbity score | 1.01 |  |
| Clashscore | 1.18 |  |
| Rotamers outliers (%) | 0.87 |  |
| Ramachandaran plot |  |  |
| Favored (%) | 96.95 |  |
| Allowed (%) | 3.05 |  |
| Outliers (%) | 0 |  |

**Supplementary Table S4.** Results of Tukey’s post hoc tests following a two-way ANOVA with repeated measures of anti-pTau antibody titre results shown in **Fig. 5c**.

|  | Day 7 | Day 14 | Day 21 | Day 28 | Day 35 |
| --- | --- | --- | --- | --- | --- |
|  | **Adjusted P Value (Summary)** | **Adjusted P Value (Summary)** | **Adjusted P Value (Summary)** | **Adjusted P Value**  **(Summary)** | **Adjusted P Value**  **(Summary)** |
| Am-S-pTau vs. Mosaic | >0.9999 (ns) | 0.9566 (ns) | 0.3339 (ns) | 0.9921 (ns) | >0.9999 (ns) |
| Am-S-pTau vs. Cocktail | >0.9999 (ns) | 0.9053 (ns) | 0.5093 (ns) | 0.9945 (ns) | >0.9999 (ns) |
| Am-S-pTau vs. Am-S-Aβ | >0.9999 (ns) | 0.8935 (ns) | 0.7018 (ns) | **<0.0001 (****)** | 0.7899 (ns) |
| Am-S-pTau vs. S-pTau + S-Aβ | >0.9999 (ns) | 0.8583 (ns) | 0.7018 (ns) | **<0.0001 (****)** | 0.7863 (ns) |
| Am-S-pTau vs. S-pTau | >0.9999 (ns) | 0.8583 (ns) | 0.7651 (ns) | **<0.0001 (****)** | 0.7863 (ns) |
| Mosaic vs. Cocktail | >0.9999 (ns) | >0.9999 (ns) | 0.9997 (ns) | 0.8903 (ns) | >0.9999 (ns) |
| Mosaic vs. Am-S-Aβ | >0.9999 (ns) | 0.9998 (ns) | **0.0102 (*)** | **<0.0001 (****)** | 0.706 (ns) |
| Mosaic vs. S-pTau + S-Aβ | >0.9999 (ns) | 0.9997 (ns) | **0.0102 (*)** | **<0.0001 (****)** | 0.7018 (ns) |
| Mosaic vs. S-pTau | >0.9999 (ns) | 0.9997 (ns) | **0.022 (*)** | **<0.0001 (****)** | 0.7018 (ns) |
| Cocktail vs. Am-S-Aβ | >0.9999 (ns) | >0.9999 (ns) | **0.0241 (*)** | **0.0007 (***)** | 0.6607 (ns) |
| Cocktail vs. S-pTau + S-Aβ | >0.9999 (ns) | >0.9999 (ns) | **0.0241 (*)** | **0.0007 (***)** | 0.6563 (ns) |
| Cocktail vs. S-pTau | >0.9999 (ns) | >0.9999 (ns) | **0.0461 (*)** | **0.0007 (***)** | 0.6563 (ns) |
| Am-S-Aβ vs. S-pTau + S-Aβ | >0.9999 (ns) | >0.9999 (ns) | >0.9999 (ns) | >0.9999 (ns) | >0.9999 (ns) |
| Am-S-Aβ vs. S-pTau | >0.9999 (ns) | >0.9999 (ns) | >0.9999 (ns) | >0.9999 (ns) | >0.9999 (ns) |
| S-pTau + S-Aβ vs. S-pTau | >0.9999 (ns) | >0.9999 (ns) | >0.9999 (ns) | >0.9999 (ns) | >0.9999 (ns) |

**Supplementary Table S5.** Results of Tukey’s post hoc tests following a two-way ANOVA with repeated measures of anti-Aβ antibody titre results shown in **Fig. 5d**.

|  | Day 7 | Day 14 | Day 21 | Day 28 | Day 35 |
| --- | --- | --- | --- | --- | --- |
|  | **Adjusted P Value (Summary)** | **Adjusted P Value (Summary)** | **Adjusted P Value (Summary)** | **Adjusted P Value**  **(Summary)** | **Adjusted P Value**  **(Summary)** |
| Am-S-Aβ vs. Mosaic | >0.9999 (ns) | >0.9999 (ns) | **0.0361 (*)** | 0.8237 (ns) | 0.9798 (ns) |
| Am-S-Aβ vs. Cocktail | >0.9999 (ns) | >0.9999 (ns) | 0.0838 (ns) | **<0.0001 (****)** | 0.9632 (ns) |
| Am-S-Aβ vs. Am-S-pTau | >0.9999 (ns) | >0.9999 (ns) | **0.0277 (*)** | **<0.0001 (****)** | 0.9279 (ns) |
| Am-S-Aβ vs. S-pTau + S-Aβ | >0.9999 (ns) | >0.9999 (ns) | **0.0277 (*)** | **<0.0001 (****)** | 0.9025 (ns) |
| Am-S-Aβ vs. S-Aβ | >0.9999 (ns) | >0.9999 (ns) | **0.0277 (*)** | **<0.0001 (****)** | 0.9025 (ns) |
| Mosaic vs. Cocktail | >0.9999 (ns) | >0.9999 (ns) | 0.9995 (ns) | **0.0010 (**)** | >0.9999 (ns) |
| Mosaic vs. Am-S-pTau | >0.9999 (ns) | >0.9999 (ns) | >0.9999 (ns) | **<0.0001 (****)** | 0.9999 (ns) |
| Mosaic vs. S-pTau + S-Aβ | >0.9999 (ns) | >0.9999 (ns) | >0.9999 (ns) | **<0.0001 (****)** | 0.9995 (ns) |
| Mosaic vs. S-Aβ | >0.9999 (ns) | >0.9999 (ns) | >0.9999 (ns) | **<0.0001 (****)** | 0.9995 (ns) |
| Cocktail vs. Am-S-pTau | >0.9999 (ns) | >0.9999 (ns) | 0.9982 (ns) | 0.6503 (ns) | >0.9999 (ns) |
| Cocktail vs. S-pTau + S-Aβ | >0.9999 (ns) | >0.9999 (ns) | 0.9982 (ns) | 0.6503 (ns) | >0.9999 (ns) |
| Cocktail vs. S-Aβ | >0.9999 (ns) | >0.9999 (ns) | 0.9982 (ns) | 0.6503 (ns) | >0.9999 (ns) |
| Am-S-pTau vs. S-pTau + S-Aβ | >0.9999 (ns) | >0.9999 (ns) | >0.9999 (ns) | >0.9999 (ns) | >0.9999 (ns) |
| Am-S-pTau vs. S-Aβ | >0.9999 (ns) | >0.9999 (ns) | >0.9999 (ns) | >0.9999 (ns) | >0.9999 (ns) |
| S-pTau + S-Aβ vs. S-Aβ | >0.9999 (ns) | >0.9999 (ns) | >0.9999 (ns) | >0.9999 (ns) | >0.9999 (ns) |

**Supplementary Methods**

***Cargo-loaded AmEnc nanocage production and characterisation***

All coding sequences were codon-optimised for expression in *E. coli* and synthesised as gBlock Gene Fragments (Integrated DNA Technologies). The encapsulin *from A. metalliredigens* (AmEnc; WP_012065385.1) was synthesised with flanking NcoI and BamHI restriction sites. The native ferritin-like protein (FLP; WP_041721970.1), containing its C-terminal encapsulation signal (Esig) peptide that mediates selective cargo recruitment into the assembling AmEnc shell, was synthesised with flanking NdeI and BglII restriction sites.

AmEnc was cloned into pACYC-Duet-1 (Novagen, Merck) and FLP into pETDuet-1 (Novagen, Merck). Both plasmids were co-transformed into *E. coli* BL21(DE3) cells (New England Biolabs, USA) for protein co-expression. Transformants were cultured aerobically at 37 °C in LB medium supplemented with carbenicillin (100 µg/mL) and chloramphenicol (50 µg/mL). Protein co-expression was induced at OD600 0.5–0.6 by addition of 0.4 mM isopropyl-β-D-thiogalactopyranoside and cultivated for 4 h at 37 °C. Cells were harvested by centrifugation (6,000 *g*, 10 min, 4 °C) and pellets stored at −30 °C until purification.

To assess iron sequestration by FLP-loaded AmEnc, cultures were grown aerobically at 37 °C in LB medium containing the appropriate antibiotics. At OD600 ~0.7–0.8, cultures were diluted with 0.4 vessel volume (v/v) of fresh medium containing 0.4 mM IPTG. After 1 h, ferrous ammonium sulphate was added to a final concentration of 4 mM, and cultures were incubated for a further 3 h at 37 °C. Cells were harvested by centrifugation (8,000 *g*, 10 min, 4 °C) and stored at −30 °C.

Purification of FLP-loaded AmEnc nanocages (with and without iron sequestration) was performed as described in the main manuscript, including heat treatment, PEG–NaCl precipitation, size exclusion chromatography, and anion-exchange chromatography. For iron-supplemented fed-batch samples, the PEG–NaCl precipitation step was omitted. All subsequent chromatographic and concentration steps were identical to those described for empty AmEnc.

Cryo-electron microscopy (cryo-EM) imaging, image processing, and three-dimensional reconstruction of FLP-loaded Am-Enc were conducted using the same procedures described for the Am-Enc structure in the main manuscript.

***Serum inflammatory cytokines***

A Cytometric Bead Array (BD Biosciences) was used to determine the concentration of pro-inflammatory cytokines in sera collected from vaccinated mice at the experimental end point. Specifically, the BD^TM^ CBA Mouse Master buffer kit, and TNF and IL-6 Flex kits were used according to the manufacturer’s protocol. Data were analysed using the BD CBA analysis software.
